# Targeting the IRE1α-XBP1 signaling axis impairs tumor growth and promotes myogenic differentiation in rhabdomyosarcoma

**DOI:** 10.1101/2025.11.10.687677

**Authors:** Anh Tuan Vuong, Aniket S Joshi, Phuong T. Ho, Meiricris Tomaz da Silva, Bin Guo, Meghana V. Trivedi, Ashok Kumar

**Author notes:** **Corresponding author:** Ashok Kumar, Ph.D., Institute of Muscle Biology and Cachexia, Department of Pharmacological and Pharmaceutical Sciences, Health Building 2, Room 5012, College of Pharmacy, University of Houston, Houston, TX 77204-1217. **Competing interest.** The authors declare no competing interests.

## Abstract

Rhabdomyosarcoma (RMS) is a pediatric soft-tissue sarcoma arising from mesenchymal progenitors with skeletal muscle features. The unfolded protein response (UPR) maintains proteostasis during endoplasmic reticulum stress, with the IRE1α-XBP1 axis representing a key signaling branch. Here, we demonstrate that components of this pathway are significantly upregulated in RMS cell lines and primary tumors. Genetic or pharmacological inhibition of IRE1α or spliced XBP1 (sXBP1) suppresses cell proliferation, promotes terminal myogenic differentiation, and enhances vincristine-induced cytotoxicity in RMS cells. Silencing of sXBP1 further reduces the cancer stem-like cell population and impairs migration and invasion. Mechanistically, IRE1α-XBP1 signaling promotes RMS progression through sXBP1-dependent upregulation of BMPR1A and subsequent activation of BMP-SMAD1 signaling. Consistently, inducible knockdown of sXBP1 or pharmacological inhibition of IRE1α endonuclease activity significantly attenuates xenograft RMS growth. Collectively, these findings identify the IRE1α-XBP1 axis as a critical regulator of RMS growth, differentiation, and chemoresistance, and support its therapeutic targeting in RMS.

## INTRODUCTION

Rhabdomyosarcoma (RMS) is a highly malignant soft tissue sarcoma that can develop throughout life but occurs more frequently in young children ^1-3^. RMS is classified into two major subtypes: embryonal rhabdomyosarcoma (ERMS) and alveolar rhabdomyosarcoma (ARMS), each with distinct histological and genetic features. ERMS often arises in the head, neck, or genitourinary tract and commonly harbors genetic alterations such as mutations in the RAS pathway or loss of heterozygosity at chromosome 11p15. Histologically, ERMS resembles immature skeletal muscle cells. ARMS is characterized by a distinctive alveolar growth pattern. ARMS commonly occurs in the extremities, trunk, or head and neck region and is associated with specific chromosomal translocations, such as PAX3::FOXO1 or PAX7::FOXO1, which generate fusion proteins that drive tumor development. For RMS patients with distant metastases, the survival rate is only about 20%-30% ^4,5^. RMS exhibits myogenic characteristics, including the expression of myogenic regulatory factors (MRFs) and muscle structural proteins. However, RMS cells fail to exit the cell cycle and undergo terminal differentiation ^2,6,7^. Therefore, agents that inhibit proliferation and promote myogenic differentiation hold promises as novel therapeutics for RMS.

Disruption of proteostasis is a hallmark of nearly all cancer types ^8^. In mammalian cells, proteostasis is maintained by several stress-responsive signaling pathways, with the unfolded protein response (UPR) of the endoplasmic reticulum (ER) being the most prominent ^9^. The ER plays a central role in the synthesis, folding, and maturation of approximately one-third of all cellular proteins. Various physiological and pathological conditions can impair protein folding, leading to the accumulation of misfolded or unfolded proteins in the ER lumen, a state referred to as ER stress. During tumor initiation, progression, and metastasis, cancer cells are exposed to numerous intrinsic and extrinsic stressors that induce ER stress. These include the loss of tumor suppressor genes, oncogene activation, oxidative stress, inflammation, hypoxia, lactic acidosis, and nutrient deprivation (e.g., glucose and amino acids) ^9,10^. In addition, conventional cancer therapies such as chemotherapy and radiotherapy can further disrupt proteostasis and exacerbate ER stress in tumor cells.

ER stress activates the UPR, which is orchestrated by the cytoplasmic portion of three ER-resident transmembrane sensors, namely inositol-requiring enzyme 1α (IRE1α), protein kinase R-like ER kinase (PERK), and activating transcription factor 6 (ATF6) ^9,11,12^. The primary function of the UPR is to restore ER homeostasis by enhancing protein folding capacity, promoting the degradation of misfolded proteins via ER-associated degradation (ERAD), and modulating protein synthesis and secretion. However, when ER stress persists and exceeds the cell’s adaptive capacity, the UPR shifts from a protective role to promoting apoptosis ^9,11,12^. Indeed, available literature suggests that cancer cells activate the UPR to alleviate ER stress caused by excessive protein-folding requirements, mutations, aneuploidy, and metabolically restrictive tumor microenvironment ^9,10^. Nevertheless, the specific roles and regulatory mechanisms of the individual UPR branches in different types of cancer remain incompletely understood.

The IRE1α is the most evolutionarily conserved ER sensor ^13^. The cytoplasmic region of IRE1α contains both a serine/threonine kinase and a tandem endoribonuclease (RNase) activity. In conditions of ER stress, IRE1α monomers dimerize and undergo autophosphorylation through their kinase domain, leading to allosteric activation of its endoribonuclease domain. The activated RNase cleaves a 26-nucleotide intronic region of X-box protein 1 (XBP1) mRNA, causing a frameshift and translation of its spliced isoform, known as spliced XBP1 (sXBP1). The sXBP1 protein is an active transcription factor that regulates the gene expression of multiple molecules involved in protein folding and secretion and ERAD pathway ^13,14^. Activated IRE1α can also cleave and degrade select mRNAs by a process termed as regulated IRE1α-dependent decay (RIDD), thereby repressing protein synthesis ^15,16^. Several recent studies have shown that the IRE1α/XBP1 signaling causes tumorigenesis and progression of several malignancies, such as multiple myeloma ^17,18^, chronic lymphocytic leukemia ^19^, breast cancer ^20-22^, and colon cancer ^23,24^. In addition to its enzymatic activity, IRE1α can also act as a scaffold protein to repress the activation of p53 protein through mechanisms potentially involving protein-protein interactions, enabling cell cycle progression in colorectal cancer and multiple myeloma ^25^.

Recent studies from our group and others have shown that the IRE1α/XBP1 signaling pathway plays a crucial role in skeletal muscle regeneration following both acute and chronic injury ^26-28^. Additionally, the IRE1α/XBP1 signaling promotes myoblast fusion during myogenic differentiation ^29^. However, its role and mechanisms of action in RMS remain unknown. In this study, we investigated how IRE1α/XBP1 signaling regulates the proliferation and differentiation of RMS cells. Our findings reveal increased expression of multiple components of the IRE1α/XBP1 pathway in RMS tumor specimens and elevated levels of phosphorylated IRE1α and spliced XBP1 (sXBP1) protein in both ERMS and ARMS cell lines. We further demonstrate that molecular or pharmacological inhibition of IRE1α/XBP1 reduces RMS cell proliferation and improves sensitivity to chemotherapeutic drug vincristine, while promoting their differentiation along the myogenic lineage. In vivo, inducible knockdown of XBP1 or pharmacological inhibition of IRE1α endonuclease activity using 4µ8C significantly inhibits tumor growth in RMS xenograft models. Furthermore, our results demonstrate that the IRE1α/XBP1 axis drives RMS progression, at least in part, through upregulating *BMPR1A* expression and enhancing BMP-Smad1 signaling.

## RESULTS

### Activation of the IRE1α/sXBP1 signaling axis in patient tumor specimens and RMS cell lines

We first analyzed how the gene expression of various components of IRE1α/XBP1 pathway is affected in human RMS samples. Analysis of a publicly available RNA-seq dataset (GSE108022) that includes normal muscle, fusion-positive RMS (FP-RMS) and fusion-negative RMS (FN-RMS) samples showed that the mRNA levels of IRE1α (gene name: *ERN1*) and XBP1 and several downstream molecules including ER chaperons, components of translocon and ERAD, and cell cycle regulators were significantly up-regulated in both FP-RMS and FN-RMS samples compared with normal muscle (**Fig. 1A**). Analysis of another publicly available microarray dataset (GSE141690) that includes 16 normal muscle and 66 RMS samples. Results showed that the mRNA levels of various components of IRE1α/XBP1 axis were significantly upregulated in RMS samples compared to normal muscle (**Fig. 1B**). We also evaluated XBP1 expression levels in different types of sarcoma cell lines using cBioPortal platform. This analysis showed that gene expression of XBP1 is increased across many types of sarcoma cell lines, including RMS (**Fig. 1C**). To further examine the involvement of the IRE1α/XBP1 signaling axis in RMS pathogenesis, we measured the levels of phosphorylated IRE1α (p-IRE1α), total IRE1α, unspliced XBP1 (uXBP1), and spliced XBP1 (sXBP1) protein in both ERMS and ARMS cell lines. Primary human myoblasts (HM), ERMS cell lines (RD, HTB82, RH36), and ARMS cell lines (RH30, RH41) were cultured under growth conditions, and protein levels were measured by performing western blot. We observed a marked increase in p-IRE1α, total IRE1α, and sXBP1 protein levels in RD, HTB82, RH36, and RH41 cells compared to HM. Additionally, there was a modest but statistically significant increase in the levels of p-IRE1α, total IRE1α, and sXBP1 protein in RH30 cells compared to HM. In contrast, uXBP1 levels were significantly reduced in most RMS cell lines, except in HTB82 cells, where a significant increase was observed compared with HM (**Fig. 1D-G**).

**FIGURE 1.**
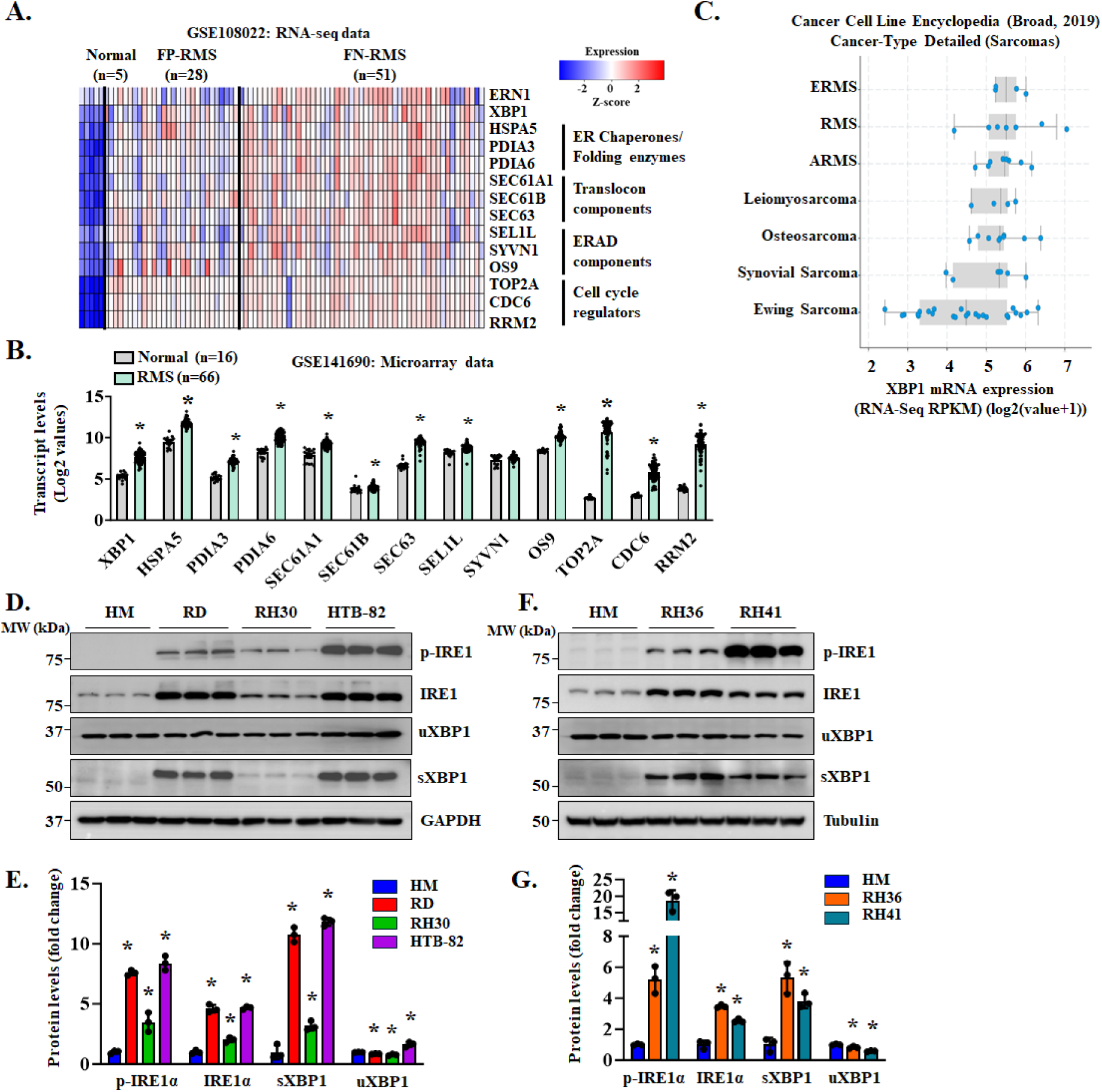
IRE1α/XBP1 signaling is activated in RMS patient tumor samples and cell lines. **(A)** Heatmap representing relative mRNA levels of IRE1α (gene name: *ERN1*), XBP1 and known XBP1-target genes in human skeletal muscle tissue (normal), fusion-positive RMS (FP-RMS), and fusion-negative RMS (FN-RMS) tumor specimens analyzed using a publicly available RNA-seq dataset (GSE108022). n=5, 28, and 51 for normal, FP-RMS and FN-RMS, respectively. Data are presented as z-scores calculated from Log2(FPKM+1) values. **(B)** Transcript levels of XBP1 and its target genes in normal human skeletal muscle tissue (n=16) and RMS tumors (n=66) analyzed using publicly available GSE141690 microarray dataset. Data are presented as mean ± SEM. *p<0.05, values significantly different from normal muscle analyzed by unpaired Student *t* test. **(C)** Box plot showing mRNA levels of XBP1 in different sarcoma cell lines analyzed using the Cancer Cell Line Encyclopedia (Broad, 2019) database on cBioPortal website. **(D)** Immunoblots and **(E)** densitometry analysis showing the levels of phosphorylated IRE1α (p-IRE1α), total IRE1α, unspliced XBP1 (uXBP1), and spliced XBP1 (sXBP1) protein in human myoblasts (HM), and RD, RH30, and HTB82 cell lines. n=3 biological replicates per group. Data are presented as mean ± SD. *p<0.05, values significantly different from HM analyzed by unpaired Student *t* test. **(F)** Immunoblots and **(G)** densitometry analysis showing the expression levels of phosphorylated IRE1α (p-IRE1α), total IRE1α, and spliced XBP1 (sXBP1) protein in HM, RH36 and RH41 cell lines. n=3 biological replicates in each group. Data are presented as mean ± SD. *p<0.05, values significantly different from HM analyzed by unpaired Student *t* test.

We also examined markers of the PERK and ATF6 branches of the UPR. Phosphorylated PERK (p-PERK) was undetectable in both HM and RMS cell lines by western blot. However, total PERK protein levels were significantly elevated in all RMS cell lines compared with HM. No significant differences were observed in phosphorylated eIF2α (p-eIF2α), total eIF2α, or CHOP protein levels between HM and RMS cell lines. Although increased expression of ATF4 or ATF6 was detected in some RMS cell lines, this was not consistently observed across all lines (**Supplementary Fig. S1**). Collectively, these results demonstrate that the IRE1α/XBP1 branch of the UPR is prominently activated in ERMS and ARMS cell lines.

### The IRE1α/XBP1 axis promotes the proliferation of RMS cell lines

To understand the role of IRE1α/XBP1 signaling in RMS, we first generated lentiviral particles expressing a scrambled (SCR), IRE1α, or sXBP1 shRNA. Cultured RD cells were transduced with lentiviral particles for 48 h and stable cells were selected by treating with 2µg/ml puromycin for 3 days. The cells were then analyzed by performing western blot for IRE1α and sXBP1 protein. Results showed that the levels of both IRE1α and sXBP1 protein were significantly reduced in RD cells stably expressing IRE1α shRNA whereas the levels of sXBP1 were reduced in RD cultures expressing sXBP1 shRNA compared to control cultures expressing SCR shRNA. In contrast, silencing of IRE1α or sXBP1 did not affect the levels of uXBP1 in RD cultures (**Fig. 2A)**. To understand how IRE1α or sXBP1 regulates gene expression in RMS cells, we performed bulk RNA sequencing (RNA-Seq) of RD cells expressing SCR, IRE1α, or sXBP1 shRNA. Results showed that IRE1α knockdown resulted in the upregulation of 691 genes and downregulation of 3,633 genes, while sXBP1 knockdown resulted in the upregulation of 532 genes and downregulation of 1,305 genes compared with controls, with differential expression deemed significant at a p-value threshold of <0.05 (**Supplementary Fig. S2A, S2B)**. Pathway enrichment analysis using Metascape Annotation tool showed that upregulated genes in IRE1α or sXBP1 knockdown RD cells were associated with skeletal system development, muscle structure development, tissue morphogenesis, cytoskeletal in muscle cells, extracellular matrix organization, mitochondria organization, cell adhesion, and regulation of cell differentiation. In contrast, downregulated genes were associated with phosphorylation, EGFR signaling, mitotic cell cycle, cell proliferation, negative regulation of cellular component organization, response to growth factor, and cell division (**Fig. 2B, C**). Heatmap analysis of the DEGs further showed that the mRNA levels of various molecules involved in cell division and proliferation were significantly downregulated in IRE1α or sXBP1 knockdown RD cultures compared with control cultures expressing SCR shRNA (**Supplementary Fig. S2C**).

**FIGURE 2.**
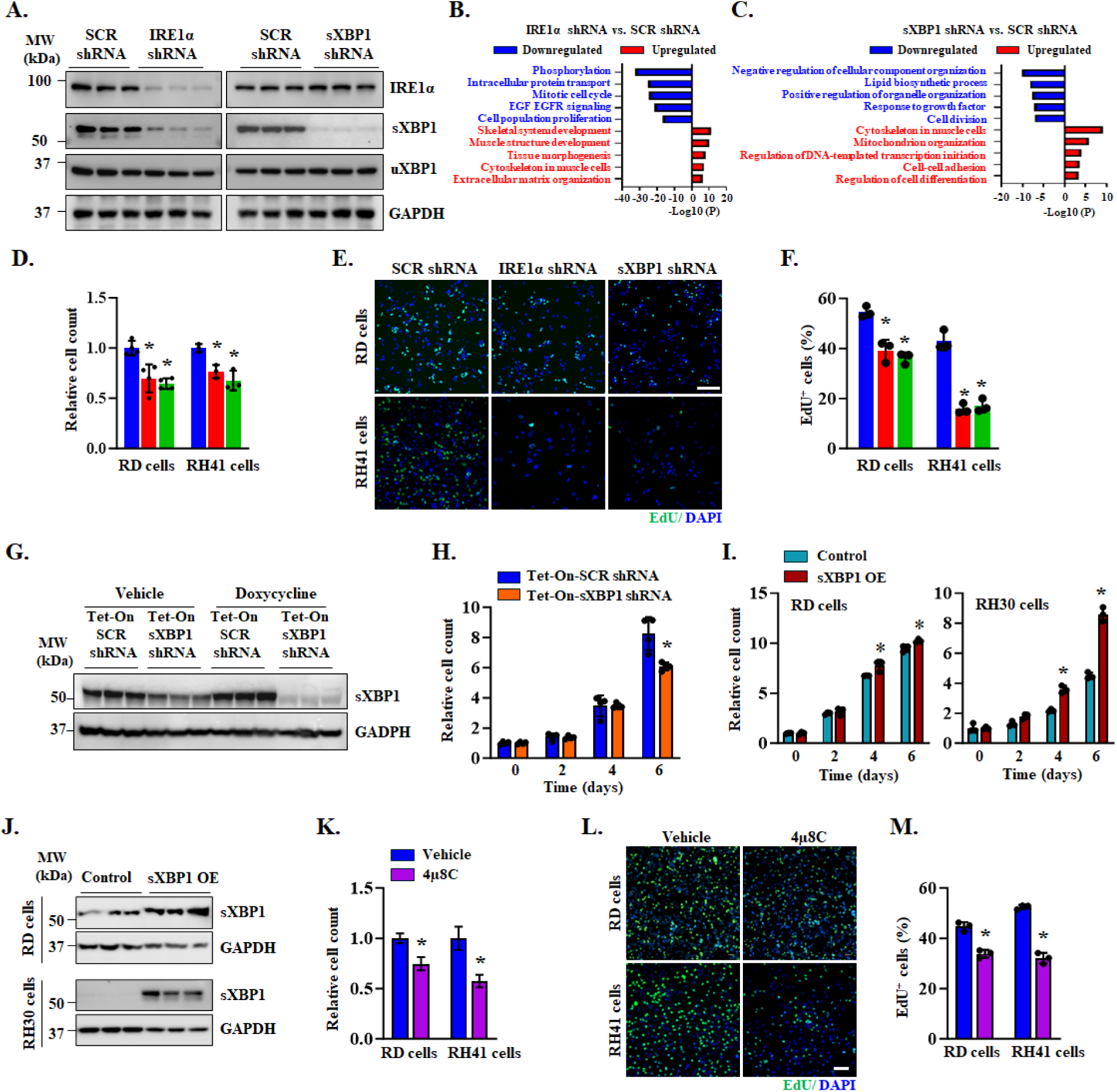
Inhibition of IRE1**α**/XBP1 signaling reduces proliferation of RMS cell lines. **(A)** Immunoblots showing the levels of IRE1α and spliced XBP1 (sXBP1) protein in scrambled (SCR), sXBP1 or IRE1α shRNA expressing RD cultures. n=3 biological replicates per group. Data are presented as mean ± SD. *p<0.05, values significantly different from SCR cultures analyzed by unpaired Student *t* test. **(B, C)** Bulk RNA-seq analysis of RD cells stably expressing SCR, IRE1α or XBP1 shRNA was performed followed by identifying differentially expressed genes (DEGs) and pathway enrichment analysis using Metascape Annotation Tool. Bar plots show enriched pathways associated with upregulated and downregulated genes in RD cells expressing **(B)** IRE1α shRNA and **(C)** XBP1 shRNA compared with those expressing SCR shRNA. **(D)** Relative cell count analyzed after 5 days of culturing SCR, IRE1α or sXBP1 shRNA expressing RD and RH41 cells. n=3 biological replicates per group. Data are presented as mean ± SD. *p<0.05, values significantly different from SCR cultures analyzed by unpaired Student *t* test. **(E, F)** RD and RH41 cells stably expressing SCR, IRE1α, or XBP1 shRNA were seeded at equal densities and incubated in growth media for 48 h, followed EdU incorporation for 2 h. **(E)** Representative photomicrographs after EdU detection. Scale bar, 100µm. **(F)** Quantification of proportion of EdU^+^ cells in IRE1α- or XBP1-knockdown RD and RH41 cultures compared with corresponding control cultures. n=3 biological replicates per group. Data are presented as mean ± SD. *p<0.05, values significantly different from SCR cultures analyzed by unpaired Student *t* test. **(G)** Immunoblots showing levels of spliced XBP1 (sXBP1) protein in RD cells expressing inducible (Tet-On) SCR or XBP1 shRNA after 72 h treatment with doxycycline or vehicle alone. n=3 biological replicates per group. **(H)** Relative number of RD cells expressing Tet-On-SCR or -XBP1 shRNA at indicated time points after treatment with doxycycline or vehicle alone. n=4 biological replicates per group. Data are presented as mean ± SD. *p<0.05, values significantly different from RD cells expressing Tet-On SCR shRNA analyzed by unpaired Student *t* test. **(I)** Relative number of RD and RH30 cells stably expressing Tet-On sXBP1 cDNA at indicated time points after treatment with doxycycline (sXBP1 OE) or vehicle alone (Control). n=4 biological replicates per group. Data are presented as mean ± SD. *p<0.05, values significantly different from control cultures analyzed by unpaired Student *t* test. **(J)** Immunoblots showing levels of sXBP1 protein in control and sXBP1 OE RD and RH30 cultures. n=3 biological replicates per group. **(K)** Relative number of RD and RH41 cells after 5 days treatment with 10 μM 4μ8C or vehicle alone. n=4 per group. Data are presented as mean ± SD. *p<0.05, values significantly different from RD or RH41 cells treated with vehicle alone analyzed by unpaired Student *t* test. **(L)** Representative photomicrographs after EdU detection. Scale bar, 100 µm. **(M)** Quantification of proportion of EdU^+^ cells in RD and RH41 cultures treated with 4μ8C or vehicle alone. n=3 biological replicates per group. Data are presented as mean ± SD. *p<0.05, values significantly different from corresponding vehicle-treated cultures analyzed by unpaired Student *t* test.

We next investigated the effect of knockdown of IRE1α or sXBP1 on the proliferation of RMS cells using RD (ERMS) and RH41 (ARMS) cell lines as models. The RD and RH41 cells expressing SCR, IRE1α, or sXBP1 shRNA were plated at equal number and total number of cells was assessed on day 5. Results showed that knockdown of IRE1α or sXBP1 significantly reduced the number of RD or RH41 cells compared to corresponding control cells expressing SCR shRNA (**Fig. 2D**). To further understand the role of the IRE1α/XBP1 pathway in RMS cell proliferation, we also performed an EdU incorporation assay. RD and RH41 cells stably expressing scrambled, IRE1α, or sXBP1 shRNA were cultured at equal densities. After 48 h, the cells were labeled with EdU for 2 h, and the proportion of EdU^+^ nuclei was detected (**Fig. 2E**). We observed a significant reduction in the proportion of EdU^+^ cells in both RD and RH41 cultures expressing IRE1α or sXBP1 shRNA compared to corresponding control cells expressing SCR shRNA (**Fig. 2F**). We also measured the effect of knockdown of IRE1α or sXBP1 on the proliferation of RH30 cells which express relatively less amounts of IRE1α or sXBP1 protein compared to other RMS cell lines (**Fig. 1D, E**). There was no significant difference in the proliferation of RH30 cells expressing SCR, IRE1α or sXBP1 shRNA (**Supplementary Fig. S2D**). This may be attributed to the relatively small increase in sXBP1 levels observed in RH30 cells, which may not be sufficient to significantly affect their proliferative capacity (**Fig. 1D, E**).

We also developed Tet-On lentiviral vectors which allow inducible expression of scrambled shRNA or sXBP1 shRNA system after treatment with doxycycline (**Fig. 2G**). Using these vectors, we investigated whether inducible knockdown of sXBP1 can also reduce the proliferation of RMS cells. Stably expressing Tet-On scrambled shRNA or Tet-on sXBP1 shRNA RD cells were plated at equal number in a 96-well plate. After 24 h, the cells were treated with 1 µg/ml doxycycline to induce sXBP1 knockdown and the number of cells were counted every two days post-doxycycline treatment. Results showed that the number of RD cells expressing Tet-On sXBP1 shRNA was significantly reduced compared to control cells transduced with Tet-On scrambled shRNA on day 6 after treatment with doxycycline (**Fig. 2H**).

We next investigated the effect of overexpression of sXBP1 on the proliferation of RMS cells. For this experiment, we engineered a Tet-On lentiviral system that allows inducible expression of sXBP1 following treatment with doxycycline. RD and RH30 cells stably expressing Tet-On sXBP1 cDNA were plated at equal number. After 24 h, the cultures were treated with 2 µg/ml doxycycline or vehicle alone, and the number of cells was counted at different time points. Results showed that sXBP1 overexpression (OE) led to a small but significant increase in the proliferation of RD cells compared to control cells on day 4 and 6 after treatment with doxycycline (**Fig. 2I**). Intriguingly, overexpression of sXBP1 protein in RH30 cells led to a two-fold increase in cell number after 6 days of treatment with doxycycline (**Fig. 2I**). This differential response between RD and RH30 cells may be attributed to the basal levels of sXBP1 protein. RD cells express higher sXBP1 levels compared to RH30 cells, leading to saturation of endogenous XBP1 in RD cells. Consequently, exogenous sXBP1 overexpression in RD cells does not significantly affect their proliferation. In contrast, the lower endogenous sXBP1 levels in RH30 cells allow for a more pronounced response to exogenous overexpression, resulting in increased cell proliferation. Western blot analysis confirmed increased sXBP1 expression in both RD and RH30 cells following doxycycline treatment (**Fig. 2J**)

To understand the therapeutic potential of inhibition of IRE1α/XBP1 arm of the UPR in RMS, we next investigated whether pharmacological inhibition of IRE1α endonuclease activity using a small molecule 4μ8C ^30^ also diminishes the proliferation of RMS cells. RD and RH41 cells were treated with vehicle alone or 10 µM 4µ8C for 72 h. Results showed that the number of RD or RH41 cells was significantly reduced in 4µ8C-treated cultures compared to corresponding cultures treated with vehicle alone (**Fig. 2K**). In another experiment, we also assessed cell proliferation by EdU incorporation assay. There was a significant reduction in the proportion of EdU^+^ cells in RD and RH41 cultures treated with 4µ8C compared to those treated with vehicle alone (**Fig. 2L, M**). However, treatment with 4µ8C or knockdown of sXBP1 had no significant effect on the proliferation of HM (**Supplementary Fig. S2E-G**).

### Inhibition of IRE1α/XBP1 signaling augments the markers of apoptosis in RMS cell cultures

Prior reports suggest that the activation of UPR protects cancer cells from undergoing apoptosis ^8,10,19,21^. We next investigated whether the inhibition of IRE1α/XBP1 axis affects the survival of RMS cells. RD and RH41 cells were transduced with lentiviral particles expressing SCR shRNA or sXBP1 shRNA. After 5 days, the cells were processed for Annexin V and propidium iodide staining followed by FACS analysis. There was a small but significant increase in the number of apoptotic (i.e., Annexin V^+^) cells observed in sXBP1 shRNA-expressing RD and RH41 cells compared to their corresponding controls (**Fig. 3A, B**). We also measured the effect of knockdown of sXBP1 on the levels of cleaved PARP and cleaved caspase-3, the two well-established biochemical markers of apoptosis ^31^. Consistent with Annexin V staining results, we observed that knockdown of sXBP1 increased the levels of cleaved PARP and cleaved caspase-3 protein in RD and RH41 cells. Silencing of sXBP1 also increased the levels of cleaved PARP and cleaved caspase-3 in HTB82 and RH30 cell lines (**Fig. 3C, D**). Moreover, we found that doxycycline-inducible knockdown of sXBP1 also increased apoptosis and levels of cleaved PARP protein in Tet-On sXBP1 shRNA-expressing RD cells (**Supplementary Fig. S3**). Finally, pharmacological inhibition of IRE1α/XBP1 axis using 4µ8C augmented the proportion of apoptotic cells and increased the levels of cleaved PARP and cleaved caspase-3 protein in RD and RH41 cells (**Fig. 3E-H**).

**FIGURE 3.**
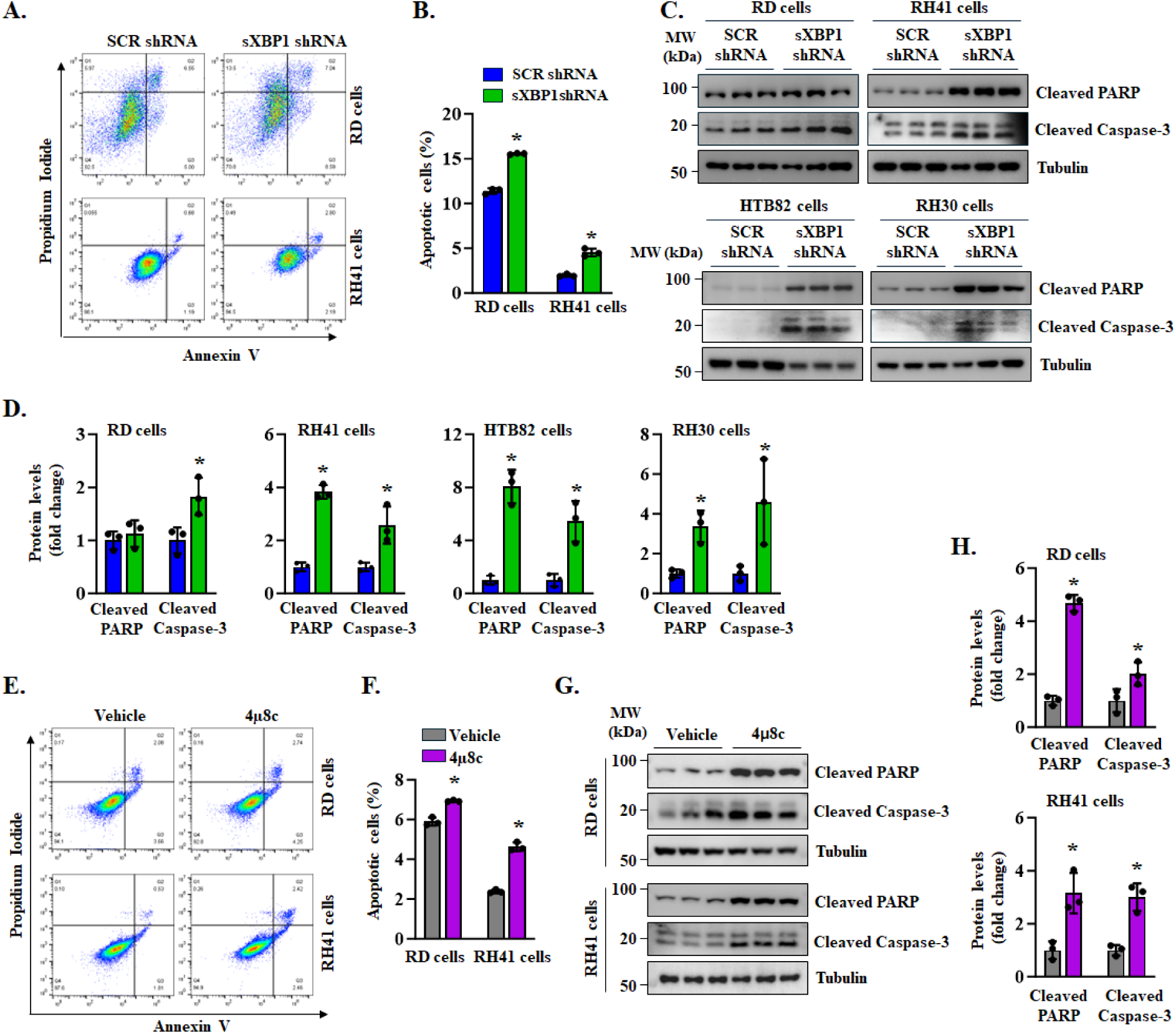
Silencing of IRE1α/XBP1 signaling promotes apoptosis in RMS cells. **(A, B)** RD and RH41 cells were transduced with lentivirus SCR or XBP1 shRNA. After 5 days, the cells were analyzed for apoptosis using propidium iodide (PI) and Annexin V staining, followed by FACS analysis. **(A)** Scatter plots and **(B)** proportion of apoptotic cells in XBP1-knockdown RD and RH41 cells compared with corresponding control cultures expressing SCR shRNA. n=3 biological replicates per group. Data are presented as mean ± SD. *p<0.05, values significantly different from corresponding cultures expressing SCR shRNA analyzed by unpaired Student *t* test. **(C)** Immunoblots and **(D)** densitometry analysis showing levels of cleaved PARP and cleaved Caspase-3 protein in SCR or XBP1 shRNA expressing RD, RH41, HTB82, and RH30 cultures. n=3 biological replicates per group. Data are presented as mean ± SD. *p<0.05, values significantly different from corresponding cultures expressing SCR shRNA analyzed by unpaired Student *t* test. **(E, F)** RD and RH41 cells were treated with 10 μM 4μ8C or vehicle alone for 3 days, followed by staining with PI and Annexin V and FACS analysis. **(E)** Scatter plots, and **(F)** quantification of proportion of apoptotic cells in vehicle- or 4µ8C-treated RD and RH41 cells. n=3 biological replicates per group. Data are presented as mean ± SD. *p<0.05, values significantly different from corresponding vehicle-treated cultures analyzed by unpaired Student *t* test. **(G)** Immunoblots, and **(H)** densitometry analysis showing levels of cleaved PARP and cleaved caspase-3 protein in RD and RH41 cultures treated with 4μ8C or vehicle alone. n=3 biological replicates per group. Data are presented as mean ± SD. *p<0.05, values significantly different from vehicle-treated cultures analyzed by unpaired Student *t* test

### Pharmacological inhibition of IRE1α/XBP1 enhances the cytotoxic effect of vincristine in RMS cells

Prior studies have shown that the activation of UPR can decrease the sensitivity of cancer cells to various chemotherapeutic agents ^32^. Vincristine is a commonly used chemotherapeutic drug for RMS ^33,34^. Therefore, we next investigated whether inhibition of IRE1α/XBP1 could enhance the anti-tumor efficacy of low-dose vincristine (2 nM for RD, RH36, RH30 and 1nM for RH41 cells) in RMS cell lines. RD, RH36, RH30, and RH41 cells were treated with vehicle alone, vincristine, 4µ8C, or a combination of vincristine and 4µ8C. After 5 days, the cells were subjected to Annexin V and PI staining followed by FACS analysis. Results showed that the proportion of apoptotic cells was significantly higher in cultures treated with the combination of vincristine and 4µ8C compared with those treated with either agent alone (**Fig. 4A, B**).

**FIGURE 4.**
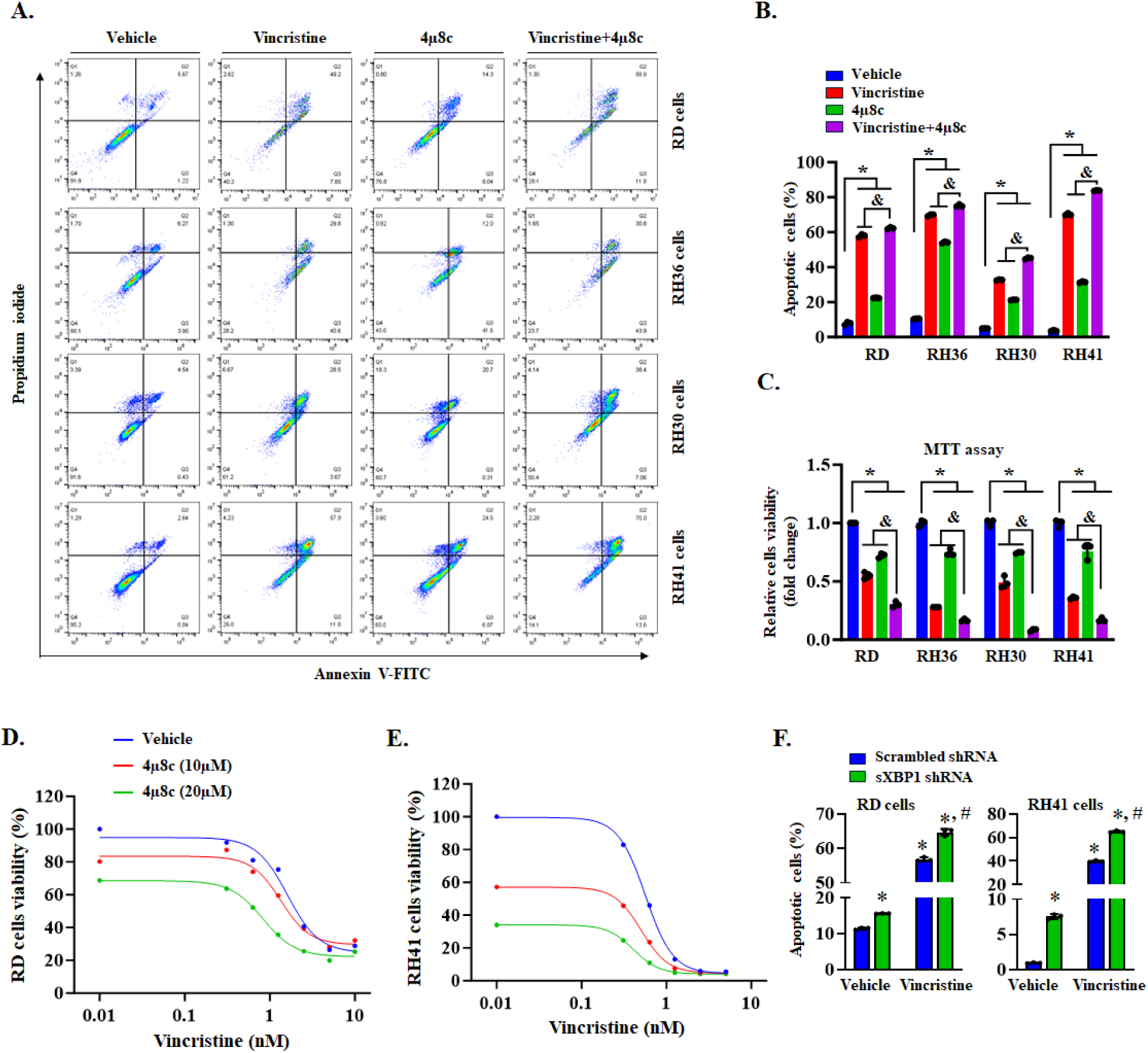
Pharmacological inhibition of IRE1α/XBP1 axis potentiates vincristine-induced cell death in RMS cultures. RD, RH36, RH30, and RH41 cells were treated with vehicle, vincristine (1nM for RH41 cells; 2nM for RD, RH36, RH30 cells), 10μM 4μ8C, or a combination of vincristine and 4μ8C for 5 days, followed by staining with propidium iodide and Annexin V and FACS analysis. **(A)** Scatter plots and **(B)** quantification of proportion of apoptotic cells in cultures treated with vehicle alone, vincristine, 4μ8C, or combination of vincristine and 4μ8C. n=3 biological replicates per group. Data are presented as mean ± SD. *p<0.05, values significantly different from corresponding vehicle-treated cultures; ^&^p<0.05, values significantly different from corresponding vincristine alone-treated cultures analyzed by unpaired Student *t* test. **(C)** Relative viability in RD, RH36, RH30, and RH41 cells after treatment with vehicle, vincristine, 4μ8C, or a combination of vincristine and 4μ8C for 5 days, assayed by performing MTT assay. n=3 biological replicates per group. Data are presented as mean ± SD. *p<0.05, values significantly different from corresponding vehicle-treated cultures; ^&^p<0.05, values significantly different from corresponding vincristine-treated cultures analyzed by unpaired Student *t* test. **(D)** RD or **(E)** RH41 cells were treated with increasing concentrations of vincristine in the presence or absence of the 4µ8C (10 µM or 20 µM), and cell viability was measured by MTT assay after 5 days. Co-treatment with 4µ8C significantly enhanced vincristine-induced growth inhibition. Combination treatment resulted in greater suppression of cell viability compared with either agent alone. **(F)** Control and XBP1 knockdown RD and RH41 cells were treated with vincristine (2 nM RD and 1 nM RH41) or vehicle alone and the percentage of apoptotic cells were determined by performing Annexin V staining and FACS analysis. n=3 biological replicates per group. Data are presented as mean ± SD. *p<0.05, values significantly different from vehicle-treated cultures expressing scrambled (SCR) shRNA. ^#^p<0.05, values significantly different from corresponding vincristine-treated cultures expressing SCR.

The MTT assay revealed that combined treatment with vincristine and 4µ8C was significantly more effective in reducing RMS cell viability than treatment with either vincristine or 4µ8C alone (**Fig. 4C**). We next sought to determine whether vincristine and 4µ8C act additively or synergistically to reduce the viability of representative ERMS and ARMS cell lines. Our initial analysis showed that the IC□□ of 4µ8C was 25.75 µM for RD cells and 7.64 µM for RH41 cells. Moreover, the IC□□ of vincristine was 1.61 nM for RD cells and 0.56 nM for RH41 cells. To evaluate the combinatorial effects of these two drugs, RD and RH41 cells were treated with increasing concentrations of vincristine in the presence or absence of 4µ8C (10 µM or 20 µM), and cell viability was measured using an MTT assay. Vincristine alone reduced cell viability in a dose-dependent manner in both RD and RH41 cells. Importantly, co-treatment with 4µ8C significantly enhanced the growth-inhibitory effects of vincristine compared with vincristine alone. In RD cells, increasing concentrations of 4µ8C (10 µM and 20 µM) markedly potentiated vincristine-induced loss of cell viability, indicating that pharmacological inhibition of IRE1α sensitizes RMS cells to vincristine treatment (**Fig. 4D**). A similar enhancement of vincristine efficacy was observed in RH41 cells (**Fig. 4E**), suggesting that this cooperative effect occurs in both embryonal and alveolar RMS subtypes.

To quantitatively assess drug interactions, combination responses were analyzed using the Bliss independence model. Bliss scores were calculated by comparing the observed inhibitory effects of the drug combination with the expected effects assuming independent drug action. The combination of vincristine and 4µ8C produced positive Bliss synergy scores across multiple dose combinations, indicating that inhibition of IRE1α signaling synergistically enhances vincristine-mediated cytotoxicity in RMS cells (**Supplementary Fig. S4A, B**).

In a separate experiment, we examined the effect of sXBP1 knockdown on vincristine-induced cytotoxicity. RD and RH41 cells were transduced with lentiviral vector expressing either scrambled (SCR) or sXBP1 shRNA for 36 h. The cells were then treated with vehicle or vincristine for 5 days, followed by Annexin V staining and FACS analysis. Knockdown of sXBP1 significantly increased vincristine-induced apoptosis in both RD and RH41 cells (**Fig. 4F, Supplementary Fig. S4C, D**). Collectively, these results demonstrate that pharmacological inhibition of IRE1α signaling using 4µ8C sensitizes RMS cells to vincristine, resulting in synergistic suppression of RMS tumor cell viability.

### Inhibition of IRE1α/XBP1 axis reduces cancer stem cells (CSCs) and epithelial-mesenchymal transition (EMT) phenotype in RD cells

Individual tumors consist of a mixed cell population that differ in function, morphology, and molecular signatures ^35-37^. Similar to other cancer types, RMS also show heterogeneity in nature, consists of a small subset of CSCs, which may be responsible for tumorigenesis, chemoresistance, and recurrence of the disease ^36-42^. CSCs of RMS express many stemness-related molecules such as Sox2, Sox9, KLF4, Oct4, and c-Myc ^39,43^. They also express CD133 on their cell surface ^40,41,43,44^. Our analysis of RNA-Seq dataset showed that mRNA levels of various markers of stemness are significantly reduced in IRE1 or sXBP1 knockdown RD cultures compared with control cultures (**Fig. 5A**). Western blot analysis also showed that the protein levels of KLF4 and Sox9 are significantly reduced in sXBP1 shRNA expressing RD cells compared to control cells expressing SCR shRNA (**Fig. 5B, C**). By performing FACS analysis, we also evaluated the effect of knockdown of sXBP1 on the proportion of CD133-expressing cells. Results showed a significant reduction in the proportion of CD133^+^ cells in sXBP1 knockdown cultures compared with control cultures (**Fig. 5D, E**).

**FIGURE 5.**
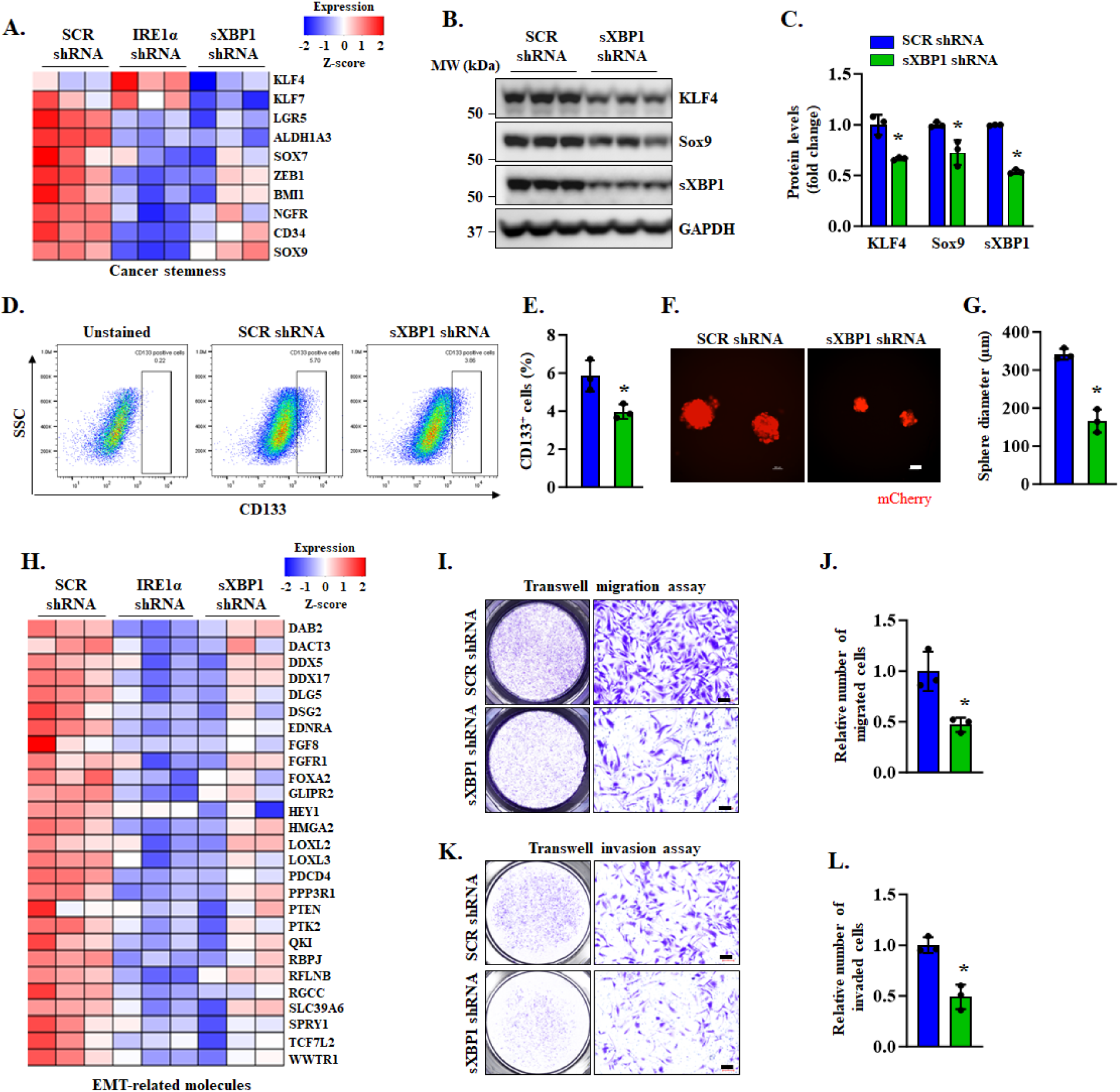
Inhibition of IRE1α/XBP1 axis represses stemness and EMT phenotype in RD cultures. **(A)** Heatmap represents relative mRNA levels of various cancer stemness-related molecules in RD cultures stably expressing scrambled (SCR), IRE1α, or XBP1 shRNA analyzed from RNA-seq dataset. **(B)** Immunoblots, and **(C)** densitometry analysis of levels of KLF4, Sox9 and sXBP1 protein in RD cultures stably expressing XBP1 shRNA compared with control cultures expressing SCR shRNA. n=3 biological replicates per group. Data are presented as mean ± SD. per group. Data are presented as mean ± SD. *p<0.05, values significantly different from SCR shRNA-expressing RD cultures analyzed by unpaired Student *t* test. **(D)** Scatter plot, and **(E)** quantification of CD133^+^ cells in RD cultures stably expressing SCR or XBP1 shRNA after performing FACS analysis. n=3 biological replicates per group. Data are presented as mean ± SD. per group. Data are presented as mean ± SD. *p<0.05, values significantly different from SCR shRNA-expressing RD cultures analyzed by unpaired Student *t* test. Data are presented as mean ± SD. *p<0.05, values significantly different from SCR shRNA-expressing RD cultures analyzed by unpaired Student *t* test. **(F)** Representative images of spheres. Scale bar, 100μm. **(G)** Quantification of average diameter of RD spheres stably expressing SCR or XBP1 shRNA. n=3 biological replicates per group. Data are presented as mean ± SD. per group. Data are presented as mean ± SD. *p<0.05, values significantly different from SCR shRNA-expressing RD cultures analyzed by unpaired Student *t* test. **(H)** Heatmap generated from RNA-seq dataset analysis shows relative transcript levels of various EMT-related genes in RD cultures stably expressing SCR, IRE1α, or XBP1 shRNA. **(I)** Representative images of migration assay. Scale bar, 100μm. **(J)** Quantification of number of migrated control and XBP1 knockdown RD cells. n=3 biological replicates per group. Data are presented as mean ± SD. *p<0.05, values significantly different from SCR shRNA-expressing RD cultures analyzed by unpaired Student *t* test. **(K)** Representative images of invasion assay. Scale bar, 100μm. **(L)** Quantification of number of invaded cells in control and XBP1 knockdown RD cultures. n=3 biological replicates per group. Data are presented as mean ± SD. *p<0.05, values significantly different from SCR shRNA-expressing RD cultures analyzed by unpaired Student *t* test.

To further elucidate the role of IRE1α/XBP1 signaling in RMS, we turned to a model system in which RMS cells are cultured in stem cell medium in suspension leading to formation of spheres, termed “rhabdospheres". This microenvironment allows only cells with stem-like properties to proliferate and form three-dimensional spheres in suspension culture ^41,43^. RD cells stably expressing scrambled or sXBP1 shRNA were cultured under non-adherent conditions in serum-free media supplemented with B27, epidermal growth factor (EGF), and basic fibroblast growth factor (bFGF) in a DMEM/F12 base. After 2 weeks of culturing, the images were captured, and diameter of the spheres was measured. Results showed that silencing of sXBP1 resulted in significantly smaller spheres compared to corresponding control cells suggesting that IRE1α/XBP1 signaling promotes the self-renewal and proliferation of CSCs in RD cell line (**Fig. 5F, G**).

Our analysis of RNA-Seq dataset also showed that the gene expression of many molecules related to EMT phenotype was significantly downregulated in RD cells following knockdown of IRE1α or sXBP1 (**Fig. 5H**). To test whether IRE1α/XBP1 axis has any role in EMT phenotypes, we studied the effect of sXBP1 silencing on the invasive and migratory capabilities of RD cells. Interestingly, stable knockdown of sXBP1 considerably reduced the migratory capacity of RD cells (**Fig. 5I, J**). Similarly, knockdown of sXBP1 led to a marked reduction in the invasive potential of RD cells (**Fig. 5K, L**) suggesting that silencing of sXBP1 diminishes the migratory and invasive capacity of RMS cells.

### Inhibition of IRE1α/XBP1 signaling augments myogenic differentiation in RMS cells

RMS is a malignant tumor of mesenchymal origin characterized by the proliferation of undifferentiated skeletal muscle cells ^2^. To determine whether activation of the IRE1α/XBP1 signaling pathway influences RMS cell differentiation, we first analyzed our RNA-seq dataset. Notably, gene expression of numerous molecules involved in muscle differentiation, structural development, and cytoskeleton organization was significantly upregulated in IRE1α- or sXBP1-knockdown RD cells compared with control cells (**Fig. 6A**), consistent with the pathway analysis results (**Fig. 2C**). To validate these transcriptomic findings, we examined the effects of IRE1α or sXBP1 knockdown on the expression of terminal differentiation markers such as myosin heavy chain (MyHC). The proportion of MyHC^+^ cells was markedly increased in RD cultures expressing IRE1α or sXBP1 shRNA compared with control cultures (**Fig. 6B, C**). Western blot analysis further confirmed elevated MyHC protein levels in IRE1α- and sXBP1-depleted RD cells (**Fig. 6D, E**). Similarly, shRNA-mediated knockdown of IRE1α or sXBP1 significantly enhanced the expression of MyHC and myogenin in RH41 cells (**Fig. 6F, G**). We also studied the effect of knockdown of sXBP1 on the differentiation of human myoblast (HM). However, there was no significant difference in the differentiation index between control and sXBP1 knockdown HM incubated in growth medium (**Supplementary Fig. S2H, I**).

**FIGURE 6.**
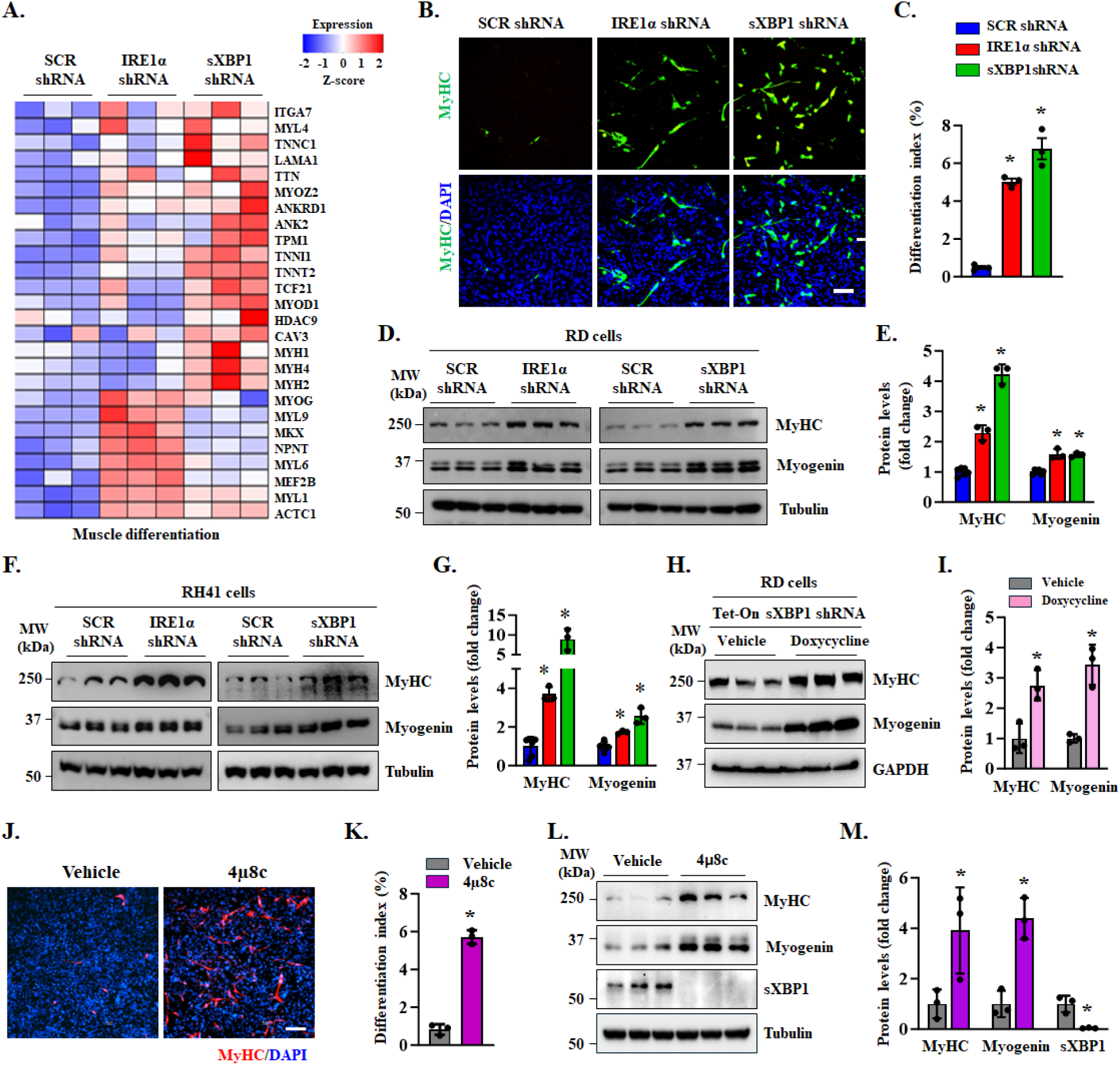
Silencing of IRE1α/XBP1 signaling axis promotes differentiation in RMS cell lines. **(A)** Heatmap generated from RNA-Seq dataset show relative mRNA levels of various muscle differentiation-related molecules in RD cultures stably expressing scrambled (SCR), IRE1α, or sXBP1 shRNA. **(B)** RD cells stably expressing SCR, IRE1α, or sXBP1 shRNA were plated at equal density and analyzed at 5 days by immunostaining for myosin heavy chain (MyHC) protein. DAPI was used to counterstain nuclei. Representative photomicrographs of MyHC-stained cultures are presented here. Scale bar, 100μm. **(C)** Quantification of differentiation index in RD cultures expressing SCR, IRE1α or sXBP1 shRNA and immunostained for MyHC. n=3 biological replicates per group. Data are presented as mean ± SD. *p<0.05, values significantly different from SCR shRNA-expressing RD cultures analyzed by unpaired Student *t* test. **(D)** Immunoblots, and **(E)** densitometry analysis of levels of MyHC and myogenin protein in SCR, IRE1α or sXBP1 shRNA expressing RD cultures. n=6 (SCR shRNA) and 3 (IRE1α or sXBP1 shRNA) biological replicates per group. Data are presented as mean ± SD. *p<0.05, values significantly different from SCR shRNA-expressing RD cultures analyzed by unpaired Student *t* test. **(F)** Immunoblots, and **(G)** densitometry analysis of MyHC and myogenin protein levels in RH41 cells stably expressing SCR, IRE1α, or sXBP1shRNA. n=6 (SCR shRNA) and 3 (IRE1α or sXBP1 shRNA). Data are presented as mean ± SD. *p<0.05, values significantly different from SCR shRNA-expressing RD cultures analyzed by unpaired Student *t* test. **(H)** Immunoblots and **(I)** densitometry analysis of levels of MyHC and myogenin protein in RD cultures expressing Tet-On sXBP1 shRNA after 5 days treatment with doxycycline or vehicle alone. n=3 biological replicates per group. Data are presented as mean ± SD. *p<0.05, values significantly different from vehicle-treated RD cultures expressing Tet-On sXBP1 shRNA analyzed by unpaired Student *t* test. **(J)** RD cells were incubated in growth media for 72 h with vehicle alone or 10μM 4μ8C followed by performing anti-MyHC and DAPI staining. Representative photomicrographs of MyHC-stained cultures are presented here. Scale bar, 100μm. **(K)** Quantification of differentiation index analyzed from MyHC-stained RD cultures treated with vehicle or 4μ8C. n=3 biological replicates per group. Data are presented as mean ± SD. *p<0.05, values significantly different from vehicle-treated RD cultures analyzed by unpaired Student *t* test. **(L)** Immunoblots, and **(M)** densitometry analysis of levels of MyHC, myogenin, and sXBP1 protein in RD cultures after 72 h of treatment with 4μ8C or vehicle alone. n=3 biological replicates per group. Data are presented as mean ± SD. *p<0.05, values significantly different from vehicle-treated RD cultures analyzed by unpaired Student *t* test.

To further assess this effect, we employed an inducible sXBP1 shRNA system. RD cells transduced with Tet-On sXBP1 shRNA lentiviral particles were treated with vehicle or doxycycline, followed by western blot analysis to determine the changes in the levels of MyHC and myogenin protein. Results showed that inducible knockdown of sXBP1 led to increased protein levels of both MyHC and myogenin (**Fig. 6H, I**). We next investigated whether pharmacological inhibition of IRE1α/XBP1 signaling would similarly promote differentiation. Treatment of RD cells with the IRE1α RNase inhibitor 4µ8C significantly increased the proportion of MyHC^+^ cells compared with controls (**Fig. 6J, K**). Consistent with these findings, western blot analysis showed elevated MyHC and myogenin and reduced sXBP1 protein levels in 4µ8C-treated cultures (**Fig. 6L, M**). Together, these results demonstrate that both molecular and pharmacological inhibition of IRE1α/XBP1 signaling enhances terminal myogenic differentiation in RMS cells.

### IRE1α/XBP1 signaling regulates BMPR1A expression in RMS cells

To elucidate the mechanisms by which IRE1α/XBP1 signaling promotes proliferation and suppresses differentiation in RMS cells, we analyzed our RNA-seq dataset. Notably, knockdown of IRE1α or sXBP1 markedly reduced the expression of several components of the BMP signaling pathway, including *BMPR1A* (**Fig. 7A**). Analysis of a publicly available gene expression dataset (GSE108022) further revealed that *BMPR1A* transcript levels were significantly elevated in both FP-RMS and FN-RMS tumors compared with normal skeletal muscle (**Fig. 7B**). Previous studies indicate that BMP signaling can exert context-dependent effects, functioning as either a tumor suppressor or promoter depending on cancer type ^45^. However, its role in RMS remains unclear. To investigate whether BMP signaling is aberrantly activated in RMS, we compared the protein levels of BMPR1A, phosphorylated Smad1 (p-Smad1), and total Smad1 between RD cells and normal human myoblasts (HM). We observed a marked increase in BMPR1A, p-Smad1, and total Smad1 protein levels in RD cells relative to HM (**Fig. 7C, D**). To explore the link between IRE1α/XBP1 signaling and BMPR1A expression, we next examined the effect of sXBP1 knockdown on BMP pathway components in RD cells. Knockdown of sXBP1 significantly decreased BMPR1A and p-Smad1 protein levels without affecting total Smad1 (**Fig. 7E, F**). Similarly, pharmacological inhibition of IRE1α RNase activity using 4µ8C reduced BMPR1A and p-Smad1 levels (**Fig. 7G, H**). Conversely, overexpression of sXBP1 increased BMPR1A and p-Smad1 protein levels in both RD and RH30 cells (**Fig. 7I, J**). Since sXBP1 functions as a transcription factor, we hypothesized that it may directly regulate *BMPR1A* transcription by binding to its regulatory region. Genome analysis using the UCSC Genome Browser identified a putative sXBP1-binding site within the enhancer region of the *BMPR1A* gene (**Fig. 7K**). To validate this prediction, we performed chromatin immunoprecipitation (ChIP) assays followed by semi-quantitative and quantitative PCR (qPCR) using primers spanning the predicted binding regions (100-200bp). ChIP results demonstrated significant enrichment of sXBP1 at the *BMPR1A* enhancer region in RD cells. The ChIP experiment was validated by performing semi-quantitative PCR for positive control RPL30 using Histone H3 antibody (**Fig. 7L**). Quantitative analysis by qPCR further showed a ∼15-fold increase in sXBP1 binding to the *BMPR1A* enhancer compared to IgG control (**Fig. 7M**). Collectively, these findings indicate that sXBP1 directly binds to and transcriptionally regulates *BMPR1A* expression, thereby modulating BMP signaling activity in RMS cells.

**FIGURE 7.**
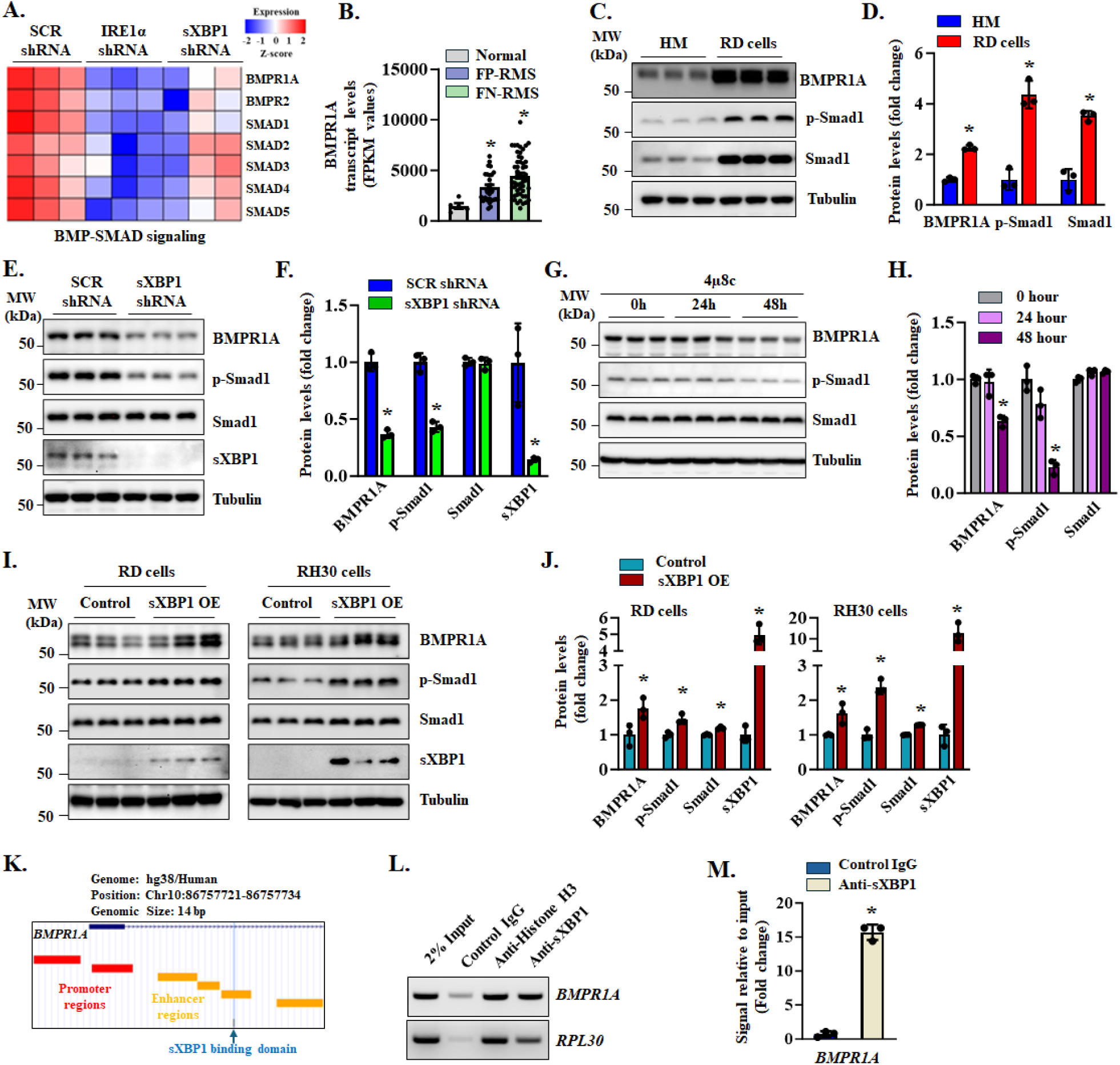
IRE1α/XBP1 signaling regulates BMPR1A expression in RMS cells. **(A)** Heatmap generated from RNA-seq dataset show relative mRNA levels of multiple BMP-SMAD signaling-related molecules in RD cultures stably expressing scrambled (SCR), IRE1α, or sXBP1 shRNA. **(B)** Bar plot shows transcript levels of BMPR1A in human skeletal muscle tissue (Normal), and fusion positive and fusion negative RMS (i.e., FP-RMS and FN-RMS, respectively) patient tumor specimens obtained by analyzing a publicly available RNA-seq database (GSE108022). **(C)** Immunoblots and **(D)** densitometry analysis showing the levels of BMPR1A, phosphorylated Smad1 (p-Smad1), and total Smad1 protein in human myoblasts (HM) and RD cultures. n=3 biological replicates per group. Data are presented as mean ± SD. *p<0.05, values significantly different from HM analyzed by unpaired Student *t* test. **(E)** Immunoblots and **(F)** densitometry analysis of levels of BMPR1A, p-Smad1, total Smad1, and sXBP1 protein in RD cultures stably expressing SCR or sXBP1 shRNA. n=3 biological replicates per group. Data are presented as mean ± SD. *p<0.05, values significantly different from SCR shRNA-expressing RD cultures analyzed by unpaired Student *t* test. **(G)** Immunoblots and **(H)** densitometry analysis of levels of BMPR1A, p-Smad1, and total Smad1 protein in RD cultures after treatment with 10μM 4μ8C for indicated time points. n=3 biological replicates per group. Data are presented as mean ± SD. *p<0.05, values significantly different from untreated (0 hour) RD cultures analyzed by unpaired Student *t* test. RD and RH30 cells stably expressing Tet-On sXBP1 cDNA were treated with doxycycline (sXBP1 OE) or vehicle alone (Control) for 5 days. **(I)** Immunoblots and **(J)** densitometry analysis of levels of BMPR1A, p-Smad1, total Smad1, and sXBP1 protein in control and sXBP1 OE RD and RH30 cultures. n=3 biological replicates per group. Data are presented as mean ± SD. *p<0.05, values significantly different from control cultures analyzed by unpaired Student *t* test. **(K)** Schematic representation of the promoter/enhancer regions flanking the transcription start site of *BMPR1A* gene. Chromatin immunoprecipitation (ChIP) assay was performed using RD cells, followed by semi-quantitative and quantitative PCR using primers designed to amplify ∼150 bp region containing the identified sXBP1 binding domain on the enhancer region of *BMPR1A* gene. **(L)** Agarose gel image shows the binding of sXBP1 protein to the promoter of *BMPR1A* gene. Validation of ChIP assay was performed using PCR for Histone H3 antibody binding to the promoter of *RPL30* (positive control). **(M)** qPCR analysis showing fold increase in the enrichment of sXBP1 to the enhancer region of *BMPR1A* gene. n=3 biological replicates per group. Data are presented as mean ± SD. *p<0.05, values significantly different from control IgG binding signal analyzed by unpaired Student *t* test.

### Inhibition of BMPR1A-mediated signaling reduces proliferation and promotes differentiation in RMS cells

Since the role of BMPR1A-mediated signaling in RMS remains unclear, we next generated lentiviral particles expressing BMPR1A shRNA to assess the effects of BMPR1A depletion on RMS cell proliferation and differentiation. RD cells were transduced with scrambled or BMPR1A shRNA for 72 h, followed by EdU incorporation assays. Results showed that knockdown of BMPR1A markedly reduced the proportion of EdU^+^ nuclei in RD cells, indicating a suppression of cell proliferation (**Fig. 8A, B**). Furthermore, BMPR1A knockdown significantly increased the expression of myogenic differentiation markers, including myogenin and MyHC, compared with control cultures (**Fig. 8C, D**). LDN193189 is a small molecule chemical compound that acts as a potent and selective inhibitor of BMP signaling which functions through inhibiting BMP type I receptors ^46^. To pharmacologically inhibit BMP signaling, RD cells were treated with vehicle alone or LDN193189 (0.1 µM) for 72 h followed by immunostaining for myogenin or MyHC protein. Interestingly, treatment with LDN193189 drastically increased the number of myogenin^+^ cells in RD cultures (**Fig. 8E, F**). Furthermore, treatment with LDN193189 increased the number of MyHC^+^ cells and differentiation index in RD cell cultures (**Fig. 8G, H**). Western blot analysis also showed that the levels of myogenin and MyHC protein are increased in LDN193189-treated RD cells compared to control cells (**Fig. 8 I, J**).

**FIGURE 8.**
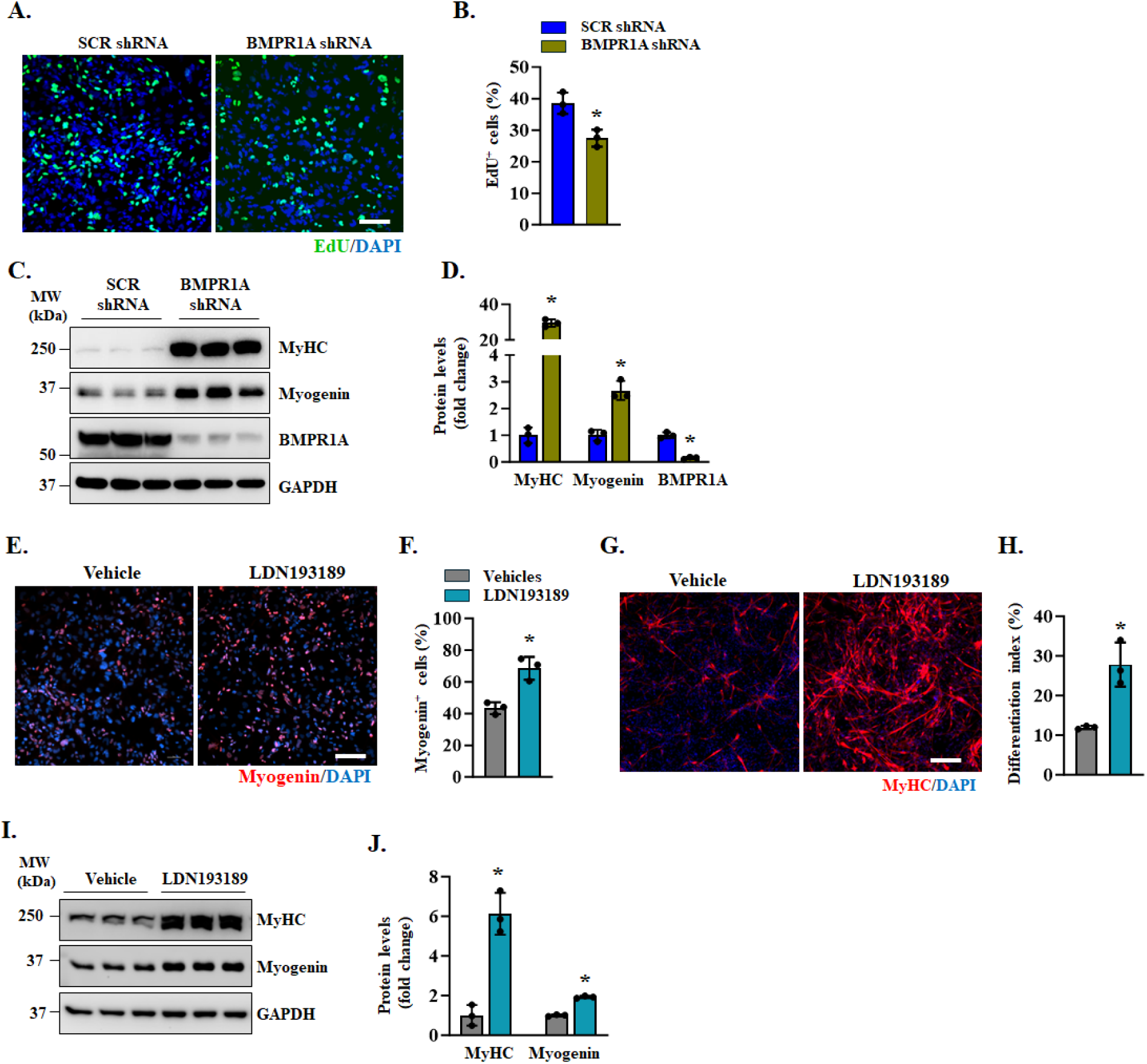
Inhibition of BMPR1A-mediated signaling suppresses proliferation and augments differentiation in RD cells. RD cells stably expressing scrambled (SCR) or BMPR1A shRNA were cultured in growth medium for 72 h, followed by EdU incorporation for 2 h. DAPI stain was used to identify nuclei. **(A)** Representative photomicrographs after EdU detection. Scale bar, 100μm. **(B)** Quantification of proportion of EdU^+^ cells in SCR or BMPR1A shRNA expressing RD cultures. **(C)** Immunoblots and **(D)** densitometry analysis of levels of MyHC, myogenin, and BMPR1A protein in SCR or BMPR1A shRNA expressing RD cultures. n=3 biological replicates per group. Data are presented as mean ± SD. *p<0.05, values significantly different from SCR shRNA expressing RD cultures analyzed by unpaired Student *t* test. RD cells were treated with 0.1μM LDN193189 or vehicle alone for 72 h followed by fixation of cells and immunostaining for myogenin or MyHC protein. Nuclei were identified by staining with DAPI. **(E)** Representative photomicrographs after immunostaining for myogenin. Scale bar, 100μm. **(F)** Quantification of proportion of myogenin^+^ cells in RD cultures treated with vehicle alone or LDN193189. **(G)** Representative photomicrographs after immunostaining for MyHC. Scale bar, 100μm. **(H)** Differentiation index in RD cultures treated with vehicle alone or LDN193189. **(I)** Immunoblots, and **(J)** densitometry analysis of levels of MyHC and myogenin protein in RD cultures treated with vehicle alone or LDN193189. n=3 biological replicates per group. Data are presented as mean ± SD. *p<0.05, values significantly different from vehicle-treated RD cultures analyzed by unpaired Student *t* test.

RNA-seq analysis showed that BMPR2 mRNA levels were also reduced along with BMPR1A. Moreover, we found that knockdown of IRE1α, but not sXBP1, significantly reduced the expression of BMPR2 in RD cells (**Fig. 7A**, **Supplementary Fig. S5A**). However, our experiments using CDD-1653, a potent and selective BMPR2 inhibitor ^47^, demonstrated that the inhibition of BMPR2 had no significant effect on the proliferation, survival, or differentiation of cultured RD cells (**Supplementary Fig. S5B-G**). Collectively, these results suggest that BMPR1A-mediated signaling supports RMS cell proliferation while restraining their terminal differentiation into skeletal muscle.

### Inhibition of IRE1α/XBP1 axis suppresses growth of RD xenografts in nude mice

To determine the role of sXBP1 in RMS tumor growth in vivo, we examined the effect of sXBP1 silencing on subcutaneous RD xenografts in nude mice. RD cells were stably transduced with Tet-On sXBP1 shRNA and firefly luciferase lentiviral particles and subsequently injected subcutaneously into the right flank of male nude mice. When the tumor size reached ∼ 150 mm^3^, the mice were randomized into two groups: one receiving doxycycline-containing chow and the other receiving standard chow. Body weight and tumor volume were monitored weekly by caliper measurements. No significant differences in body weight were observed between the two groups throughout the study (**Fig. 9A**). Notably, within two weeks of doxycycline administration, tumor volumes in the doxycycline-treated group were significantly reduced compared with controls, and this difference continued to increase through day 28 (**Fig. 9B**). In vivo bioluminescence imaging following luciferin injection confirmed markedly smaller tumors in doxycycline-treated mice after 25 days (**Fig. 9C**). At the end of the study, tumors were excised and weighed. Tumor wet weight in the doxycycline-treated group was reduced by approximately 50% relative to controls (**Fig. 9D, E**). To exclude potential effects of doxycycline alone, a parallel experiment was performed using RD cells expressing Tet-On scrambled shRNA and luciferase. No significant differences in body weight, tumor volume, or tumor wet weight were observed between doxycycline-treated and control mice in this cohort (**Supplementary Fig. S6**), indicating that doxycycline treatment per se does not affect RD tumor growth.

**FIGURE 9.**
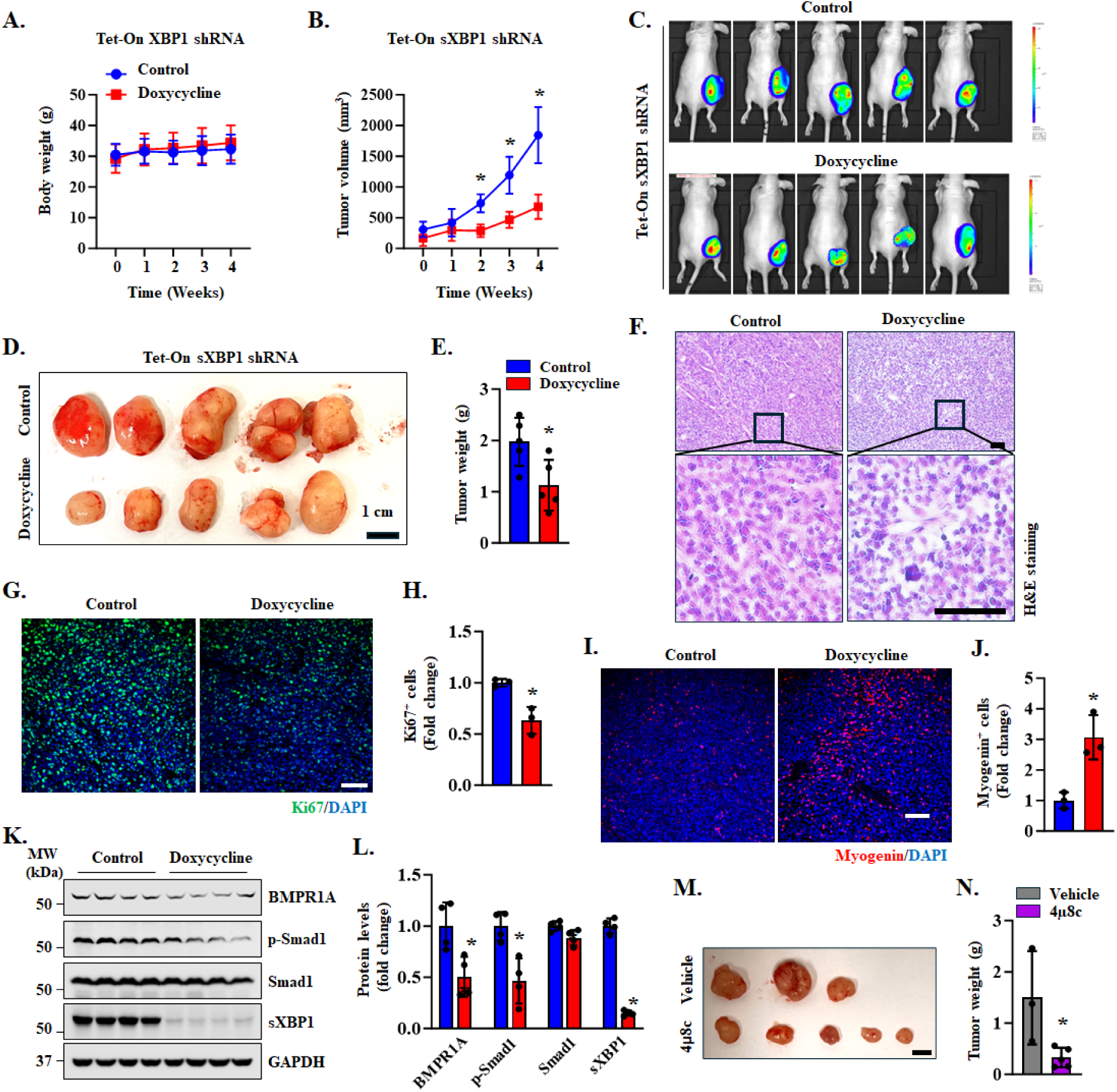
Inducible knockdown of sXBP1 suppresses tumor growth in RD xenografts. **(A)** Body weight of mice inoculated with Tet-On sXBP1 shRNA expressing RD cells fed with normal chow (control) or doxycycline-containing chow. **(B)** Average tumor volume in mice fed with normal chow or doxycycline containing chow. n=5 mice in each group. Data are presented as mean ± SD. *p<0.05, values significantly different from normal chow fed mice at indicated time points by unpaired Student *t* test. **(C)** Bioluminescence imaging showing presence RD xenograft in control and doxycycline-treated nude mice. **(D)** Images of tumors of control and doxycycline-treated group at the time of euthanizing the mice. Scale bar, 1cm. **(E)** Quantification of wet weight of tumors in control and doxycycline-treated group. n=5 mice in each group. Data are presented as mean ± SD. *p<0.05, values significantly different from control group by unpaired Student *t* test. **(F)** Representative images of tumor sections after performing H&E staining. Scale bar, 100µm. **(G)** Representative images of tumor sections after performing immunostaining for Ki67 protein. Nuclei were counterstained with DAPI. Scale bar, 100µm. **(H)** Quantification of relative number of Ki67^+^ cells in tumor samples of control and doxycycline-treated mice. n=3 mice/group. Data are presented as mean ± SD. *p<0.05, values significantly different from control group by unpaired Student *t* test. **(I)** Representative images of tumor sections after performing immunostaining for myogenin protein. Nuclei were counterstained with DAPI. Scale bar, 100µm. **(J)** Quantification of relative number of myogenin^+^ cells in tumor samples of control and doxycycline-treated mice. n=3 per group. Data are presented as mean ± SD. *p<0.05, values significantly different from normal chow fed mice analyzed unpaired Student *t* test. **(K)** Immunoblots and **(L)** densitometry analysis of levels of sXBP1, BMPR1A, p-Smad1, and total Smad1 in tumor samples of doxycycline-treated mice compared to control mice. n=4 mice per group. Data are presented as mean ± SD. *p<0.05, values significantly different from control group by unpaired Student *t* test **(M)** Images of RD tumors at the time of euthanizing the mice that were treated twice per week with 4μ8C (10 mg/kg body weight) or vehicle alone. Scale bar, 1cm. **(N)** Quantification of wet weight of tumors in vehicle- or 4μ8C-treated mice. n=3 (vehicle-treated) and 5 (4μ8C-treated) mice. Data are presented as mean ± SD. *p<0.05, values significantly different from control group by unpaired Student *t* test.

We next performed histological and biochemical analyses of tumor samples. Hematoxylin and eosin (H&E) staining of tumor sections revealed reduced tumor cell density and the presence of multinucleated cells in sXBP1 knockdown tumors, suggestive of growth arrest and myogenic differentiation (**Fig. 9F**). Immunostaining for Ki67, a proliferation marker, demonstrated a marked reduction in Ki67^+^ cells in doxycycline-treated tumors (**Fig. 9G, H**). Conversely, there was a significant increase in myogenin^+^ cells, indicating enhanced myogenic differentiation (**Fig. 9I, J**). Western blot analysis confirmed efficient sXBP1 knockdown, as evidenced by a substantial reduction in sXBP1 protein levels in tumors of doxycycline-treated mice. Additionally, BMPR1A and phosphorylated Smad1 (p-Smad1) protein levels were also significantly reduced in tumors of doxycycline-treated mice compared with controls (**Fig. 9K, L**). To complement these molecular findings, we also evaluated the effect of pharmacological inhibition of IRE1α/XBP1 signaling on RD xenograft growth. Nude mice bearing subcutaneous RD tumors (∼150 mm³) were treated with vehicle or the IRE1α RNase inhibitor 4µ8C (10 mg/kg) twice per week by intraperitoneal injection for 4 weeks. After 4 weeks of treatment, the final dose of 4µ8C was administered 12 h before euthanizing the mice. Treatment with 4µ8C significantly reduced RD tumor size in mice compared with vehicle controls (**Fig. 9M, N**). Together, these results demonstrate that both molecular and pharmacological inhibition of the IRE1α/XBP1 axis markedly suppresses RMS tumor growth in vivo.

## DISCUSSION

Despite significant therapeutic advances, the prognosis for children with aggressive or metastatic RMS remains poor, largely because the molecular mechanisms driving tumorigenesis and chemoresistance are not fully understood ^4,7,34,48^. Accumulating evidence indicates that the IRE1α/XBP1 branch of the unfolded protein response (UPR) exerts context-dependent effects in cancer, generally promoting tumor growth and therapeutic resistance through adaptation to ER stress, yet functioning as a tumor suppressor under specific conditions ^22,49-51^. However, the role and mechanisms by which the IRE1α/XBP1 signaling axis regulates RMS pathogenesis remain poorly understood.

Through integrated analyses of patient-derived tumor datasets, RMS cell lines, and xenograft models, we demonstrate that the IRE1α/XBP1 axis is constitutively activated in RMS subtypes. These findings are consistent with reports of persistent IRE1α/XBP1 activation in other malignancies, including breast, prostate, and pancreatic cancers, sarcomas, and glioblastoma ^52-55^. We further show that genetic or pharmacological inhibition of IRE1α/XBP1 signaling suppresses RMS cell proliferation and survival, promotes terminal myogenic differentiation, and enhances sensitivity to chemotherapeutic agents such as vincristine. Finally, our experiments demonstrate that sXBP1 directly regulates BMPR1A transcription, thereby linking UPR activation to BMP-Smad signaling and the maintenance of an undifferentiated, stem-like phenotype in RMS.

The activation of IRE1α/XBP1 signaling in RMS appears to be a survival mechanism that supports the high biosynthetic demands of rapidly proliferating tumor cells. Silencing of IRE1α or sXBP1 reduced proliferation of both ERMS and ARMS cell lines whereas forced expression of sXBP1 augmented the proliferation of RH30 cells. RNA-seq analysis following knockdown of IRE1α or sXBP1 further revealed downregulation of many cell cycle-related genes, particularly those involved in mitotic progression and growth factor signaling, suggesting that this pathway promotes RMS cell proliferation at the transcriptional level (**Fig. 2**). Interestingly, gene expression of many molecules involved in muscle structure development and differentiation was upregulated, suggesting that the inhibition of IRE1α/XBP1 axis mitigates the impaired differentiation in RMS (**Fig. 6**). Consistent with these transcriptional changes, our results demonstrated that pharmacologic inhibition of IRE1α RNase activity with 4μ8C reduced proliferation and survival and improved terminal differentiation in RMS cells.

Our results further demonstrate that inhibition of the IRE1α/XBP1 axis enhances the cytotoxic efficacy of vincristine in cultured RMS cell lines. The combination of vincristine and 4µ8C produced greater cytotoxic effects than either agent alone. Moreover, Bliss independence analysis yielded positive synergy scores, indicating a synergistic interaction between the two molecules (**Fig. 4, Fig. S4**). These findings are consistent with previous studies showing that activation of the UPR can protect cancer cells from chemotherapeutic stress by limiting ER stress-induced apoptosis ^32,56^. For example, constitutive IRE1α RNase activity promotes the production of pro-tumorigenic factors and confers resistance to paclitaxel in triple-negative breast cancer, whereas inhibition of IRE1α enhances paclitaxel-induced tumor regression and delays relapse ^57^. Similarly, activation of the IRE1α/XBP1 pathway promotes chemoresistance in lung cancer, partly through increased expression of multidrug resistance protein-1, which facilitates drug extrusion ^58^. Although the underlying mechanisms remain less understood, previous studies have also reported that pharmacological inhibition of UPR components reduces the survival of cultured RMS cells ^55^. Consistent with these observations, our results suggest that RMS cells are selectively dependent on IRE1α/XBP1 signaling for proliferation, survival, and chemoresistance, highlighting this pathway as a potential therapeutic target.

Recent studies have implicated the UPR in the maintenance of stemness in several malignancies, including glioblastoma, breast cancer, and hepatocellular carcinoma, where XBP1 activation enhances self-renewal and chemoresistance ^52-54^. Consistent with these observations, our results show that inhibition of the IRE1α/XBP1 pathway reduces rhabdosphere formation and suppresses the expression of genes associated with stemness and EMT (**Fig. 5**). Cancer stem cells frequently exhibit EMT-associated features such as increased plasticity, self-renewal capacity, and therapeutic resistance, and EMT-related transcriptional programs can maintain tumor cells in a dedifferentiated state ^59^. Accordingly, repression of EMT-related genes following sXBP1 knockdown suggests that the IRE1α/XBP1 pathway helps sustain a stem-like phenotype in RMS cells. Moreover, our results demonstrating that the inhibition of IRE1α/XBP1 axis promotes the expression of myogenic differentiation markers (**Fig. 6**) indicate that suppression of this pathway may shift the cellular state from stem-like toward a more differentiated phenotype.

Although the precise mechanisms remain unclear, sXBP1 may promote stemness by directly regulating genes involved in stem cell maintenance or indirectly by controlling cellular processes that support adaptation to stress, including protein folding, redox balance, and lipid biosynthesis. In addition, IRE1α signaling may interact with oncogenic pathways such as PI3K/AKT, Wnt/β-catenin, and Notch ^50,60^, thereby reinforcing stemness networks and promoting RMS cell survival under stress. Beyond stemness regulation, our RNA-seq analysis revealed downregulation of multiple EMT-associated genes following knockdown of IRE1α or sXBP1, and transwell assays confirmed that sXBP1 silencing reduced the migratory and invasive capacities of RMS cells (**Fig. 5**). Collectively, these findings suggest that IRE1α/XBP1 signaling promotes both stemness and EMT phenotypes in RMS, processes that contribute to tumor aggressiveness. Future studies will determine whether sXBP1 directly regulates genes involved in stemness and EMT programs.

One of the hallmarks of RMS is the failure of myogenic differentiation ^2,6,61^. Our results show that inhibition of the IRE1α/XBP1 signaling axis increases the expression of differentiation markers such as myogenin and MyHC in both ERMS and ARMS cells, suggesting that this pathway actively represses terminal myogenic differentiation (**Fig. 6**). During normal myogenesis, BMPR1A-Smad1 signaling pathway regulates the balance between progenitor proliferation and differentiation ^62,63^. However, persistent BMP signaling can maintain cells in a proliferative state and promote tumorigenesis ^45,64,65^. Consistent with this concept, RMS cells exhibited higher BMPR1A and p-Smad1 levels than normal human myoblasts, and both molecular and pharmacological inhibition of XBP1 reduced their expression (**Fig. 7**) Furthermore, we identified direct binding of sXBP1 to the BMPR1A enhancer, revealing a previously unrecognized IRE1α-XBP1-BMPR1A regulatory axis linking ER stress signaling with pathways controlling cell fate in RMS.

Previous studies have also implicated dysregulation of the TGF-β/Smad2/3 signaling pathway and the BMP-driven Smad1/5/8 signaling pathway in RMS pathogenesis through their effects on proliferation and differentiation ^62,66-68^. However, therapeutic targeting of these pathways may be challenging because of their essential roles in normal development, tissue repair, and immune regulation. In contrast, the IRE1α/XBP1 signaling axis primarily functions as a cellular adaptation mechanism to ER stress. Cancer cells often rely on this pathway to sustain rapid proliferation and survive under stressful tumor microenvironment conditions. Our findings indicate that inhibition of this pathway suppresses RMS growth and enhances sensitivity to vincristine, suggesting that it may represent a more tumor-selective therapeutic vulnerability. Nonetheless, given the potential crosstalk between ER stress signaling and TGF-β/BMP pathways, future studies should explore whether combinatorial targeting of these signaling networks could further improve therapeutic outcomes in RMS.

Our experiments also showed that inducible knockdown of sXBP1 markedly suppresses the growth of RD xenografts in nude mice, establishing a crucial role for XBP1 in promoting RMS tumor progression in vivo. Using a doxycycline-inducible shRNA system, we demonstrated that suppression of sXBP1 leads to a substantial reduction in tumor volume and weight. Consistent with the genetic approach, pharmacological inhibition of IRE1α RNase activity using 4µ8C produced comparable tumor-suppressive effects, further validating the functional importance of the IRE1α/XBP1 signaling axis in RMS growth. Reduced BMPR1A and p-Smad1 levels in XBP1-deficient tumors suggest that sXBP1 sustains RMS growth, at least in part, through BMP/Smad signaling. Moreover, the reduction in tumor burden following sXBP1 knockdown was accompanied by decreased proliferation and enhanced differentiation of RD cells (**Fig. 9**), consistent with our in vitro findings showing that inhibition of IRE1α/XBP1 signaling relieves the differentiation block in RMS cells (**Fig. 6**).

Although our in vivo data show that molecular or pharmacological inhibition of the IRE1α/XBP1 axis reduces RD xenograft growth, further studies are required to determine whether similar effects occur in RMS patient-derived xenograft (PDX) models. In addition, our in vitro findings demonstrate that knockdown or pharmacological inhibition of XBP1 enhances vincristine-induced cytotoxicity in multiple RMS cell lines. Future studies should therefore examine whether targeting sXBP1 also improves the therapeutic efficacy of vincristine against RMS in vivo.

In summary, our study provides initial evidence that the IRE1α/XBP1 signaling axis is a central regulator of RMS growth. Since this pathway intersects multiple oncogenic processes, including proliferation, survival, stemness, and differentiation blockade, its inhibition represents a multifaceted approach to RMS therapy.

## METHODS

### Cell culture

RD, RH30, and HTB82 rhabdomyosarcoma cell lines were obtained from the American Type Culture Collection (ATCC). RH36 and RH41 cell lines were kindly provided by the Houghton laboratory (UT Health San Antonio, Texas). RD and RH36 cells were maintained in Dulbecco’s modified Eagle medium (DMEM) supplemented with 10% fetal bovine serum (FBS) and 1% penicillin-streptomycin (P/S). RH30 and RH41 cells were cultured in RPMI 1640 medium supplemented with 10% FBS and 1% P/S. HTB82 cells were cultured in McCoy’s 5A medium supplemented with 10% FBS and 1% P/S. Human skeletal muscle myoblasts were purchased from ThermoFisher Scientific and maintained in Iscove’s Modified Dulbecco’s Medium (IMDM) supplemented with 20% FBS, 10% horse serum, and 20 ng/mL basic fibroblast growth factor. All cell lines were routinely tested and confirmed to be mycoplasma-free by using Universal Mycoplasma Detection Kit (ATCC 30-1012K, ATCC, USA).

### Generation of plasmid constructs

The target siRNA sequence for human IRE1α, XBP1, or BMPR1A mRNA was identified using BLOCK-iT™ RNAi Designer online software (Life Technologies). The shRNA oligonucleotides were synthesized to contain the sense strand of target sequences for human IRE1α (5’- GGT GGA ATG CCA CCT ACT TTG-3’), human sXBP1 (5’-GGA ATG ATC CAA TAC TGT TGC-3’), or human BMPR1A (5’-GGA GAA ACC ACA TTA GCT TCA-3’), short spacer (CTCGAG), and the reverse complement sequences followed by five thymidines as an RNA polymerase III transcriptional stop signal. Oligonucleotides were annealed and cloned into pLKO.1-mCherry-Puro with AgeI/EcoRI sites. For inducible knockdown, the XBP1 shRNA sequence was inserted into the pLKO-Tet-On plasmid (Addgene plasmid #21915). For Tet-On–inducible overexpression of sXBP1, the coding sequence of sXBP1 was subcloned from pCMV5-Flag-XBP1s (Addgene plasmid #63680) into the pLKO-Tet-On backbone.

### Lentivirus production, transduction, and stable expression

Lentiviral particles were generated following a protocol as described ^29^. Briefly, pLKO.1 or pLKO-Tet-On plasmids encoding scrambled, IRE1α, sXBP1, or BMPR1A shRNA were co-transfected with packaging plasmids psPAX2 and pMD2.G into HEK293T cells using polyethylenimine (PEI; 1 µg/mL). After 48 h, culture supernatants were collected, centrifuged at 4,500 rpm for 15 min to remove cell debris, and filtered through a 0.45-µm membrane. RMS cells were transduced with viral supernatants by centrifugation in the presence of 6 to 10 µg/ml polybrene (Sigma-Aldrich). After 24 h, the medium was replaced with fresh growth medium. To establish stable cell lines, transduced cells were selected with 2 µg/mL puromycin for 2-3 passages beginning 72 h post-transduction. For inducible knockdown or overexpression, Tet-On RMS cells were treated with 1 µg/ml doxycycline for 3-5 days prior to experiments.

### Cell proliferation assay

Cells were seeded at 1,000 cells per well in PhenoPlate™ 96-well tissue culture plates (Revvity, USA). Plates were scanned using the EnSight Multimode Plate Reader with well-imaging technology (PerkinElmer, MA), and cell counts were obtained through digital phase and bright-field imaging. To evaluate pharmacological effects, cells were treated with chemical compounds after overnight seeding, followed by regular scanning and cell counting. For experiments using the Tet-On system, cells were treated with doxycycline (1 µg/ml) after overnight seeding, with subsequent imaging and counting performed at regular intervals.

### EdU incorporation assay

DNA synthesis was assessed using the Click-iT™ Plus EdU Alexa Fluor™ 488 Imaging Kit (Invitrogen, USA). RMS cells were incubated with 1 µg/mL EdU for 2 h, fixed with 4% paraformaldehyde, and permeabilized with 0.5% Triton X-100. EdU incorporation was visualized using the Click-iT reaction with Alexa Fluor™ 488 dye, and nuclei were counterstained with DAPI. Images were acquired using a fluorescence microscope, and the percentage of EdU□ nuclei relative to total DAPI□ nuclei was quantified using NIH ImageJ software. Three to four images from each biological replicate were analyzed.

### Cell viability assay

Approximately 5 × 10³ cells were seeded in 6-well plates containing 2 ml of culture medium per well and maintained for 5 days with medium change after 48 h. Dead cells were removed by washing the wells with PBS. The adherent cells were fixed and stained with 0.25% crystal violet dye in 20% methanol and images were captured. For quantification, the bound dye was solubilized in 1 ml of 10% acetic acid per well, incubated for 15 min at room temperature with gentle shaking, diluted 1:4 with water, and absorbance was measured at 590 nm using a SpectraMax® i3x microplate reader (Molecular Devices).

### Sphere formation assay

Approximately 1 × 10³ RD cells stably transduced with scrambled or XBP1 shRNA were seeded per well in 24-well low-attachment Nunclon™ Sphera™ dishes (Thermo Fisher Scientific, USA). Each well contained 1 ml of DMEM/F12 medium supplemented with B27, epidermal growth factor (EGF, 10 ng/mL), and basic fibroblast growth factor (bFGF, 10 ng/mL). After 14 days of incubation, the images of the spheres were acquired using a microscope and the diameter of the sphere was measured.

### Immunofluorescence

RMS cells stably expressing scrambled (SCR), IRE1α, or XBP1 shRNA were cultured for 5 days. The cultures were washed with PBS, fixed with 4% paraformaldehyde (PFA) for 15 min at room temperature, and permeabilized with 0.5% Triton X-100 in PBS for 15 min. Cells were then blocked with 5% goat serum in PBS containing 0.1% Triton X-100. The anti-MyHC (clone MF20) primary antibody was diluted 1:50 in 5% goat serum and incubated with the cells overnight at 4°C. After washing with PBST, cells were incubated with goat anti-mouse secondary antibody (diluted 1:300 in 5% goat serum) for 1 h at room temperature. Nuclei were counterstained with DAPI for 15 min. Images were acquired using a fluorescence inverted microscope (Nikon Eclipse TE2000-U) equipped with a digital camera (Digital Sight DS-Fi1). The percentage of MyHC cells was quantified using ImageJ software.

### Differentiation index

Differentiation of RMS cells was quantified by measuring the differentiation index which is defined as: (Number of nuclei in MyHC^+^ cells/Total number of nuclei) × 100.

### Transwell migration and invasion assay

Transwell assays were performed using Boyden chambers (Corning Inc., Corning, NY, USA). For migration assays, 5 × 10^4^ RD or RH30 cells in 200 μl serum-free DMEM were seeded into the upper chamber, while 800 μl of DMEM containing 10% FBS was added to the lower chamber as a chemoattractant. After 24 h incubation, cells that migrated through the membrane were fixed and stained with 0.1% crystal violet. For invasion assays, the procedure was identical except that the upper surface of the membrane was pre-coated with 100 μL of 300 μg/mL Matrigel (BD Biosciences, San Jose, CA, USA). Images of crystal violet-stained cells on the lower membrane surface were captured, and signal intensity normalized to total area was quantified using ImageJ software (NIH).

### Annexin V staining and FACS analysis

Cell viability and apoptosis were assessed by Annexin V-FITC and propidium iodide (PI) staining using the Annexin V-FITC Apoptosis Detection Kit, according to the manufacturer’s instructions (Abcam). Stained cells were analyzed by flow cytometry.

### MTT assay

Cell growth and viability of control and IRE1α- or XBP1-knockdown RMS cells were evaluated using the MTT assay. Briefly, RMS cells stably expressing scrambled, IRE1α, or sXBP1 shRNA were seeded at 5 × 10^3^ cells per well in a 96-well plate and cultured for 3 days. Cells were then incubated with MTT solution (0.5 mg/mL) for 4 h at 37°C. The medium was removed, and 100 μl of DMSO was added to each well to dissolve the formazan crystals. Absorbance was measured at 570 nm using a SpectraMax® i3x microplate reader (Molecular Devices). Cell viability was expressed as the percentage relative to corresponding control cells.

### Drug combination and synergy analysis

Drug interaction between vincristine and the 4µ8C inhibitor was analyzed using the web-based platform SynergyFinder. RD and RH41 cells were treated with increasing concentrations of vincristine in the presence or absence of increasing concentrations of 4µ8C, and cell viability was determined using the MTT assay. The resulting viability values were normalized to vehicle-treated controls and imported into SynergyFinder for analysis. Drug interaction scores were calculated using the Bliss independence model, which predicts the expected combined response of two drugs assuming independent mechanisms of action. Bliss synergy scores were computed by comparing the observed inhibitory effect of the drug combination with the expected effect predicted by the model. Positive Bliss scores indicate synergistic interactions, values near zero indicate additive effects, and negative scores indicate antagonistic interactions. Synergy landscapes and dose-response interaction maps were generated using SynergyFinder to visualize regions of synergy across the drug combination matrix. The overall interaction between vincristine and 4µ8C was summarized by the average Bliss synergy score across all tested dose combinations.

### Western Blot

Cultured RMS cell lines or RD tumor samples were washed with PBS and homogenized in lysis buffer A (50 mM Tris-Cl [pH 8.0], 200 mM NaCl, 50 mM NaF, 1 mM dithiothreitol, 1 mM sodium orthovanadate, 0.3% IGEPAL, and protease inhibitors). Approximately, 40 μg protein was resolved on each lane on 8-10% SDS-PAGE gel, transferred onto a PVDF membrane, and probed using a specific primary antibody described in **Supplementary Table S1**. Bound antibodies were detected by secondary antibodies conjugated to horseradish peroxidase (Cell Signaling Technology). Signal detection was performed by an enhanced chemiluminescence detection reagent (Bio-Rad). Approximate molecular masses were determined by comparison with the migration of prestained protein standards (Bio-Rad). Uncropped gel images are presented in **Supplementary Fig. S7**.

### Chromatin immunoprecipitation (ChIP) assay

ChIP was performed using SimpleChIP enzymatic Chromatin IP kit (Cell Signaling Technology, Cat #9003) according to the manufacturer’s suggested protocol. Briefly, 6 x 10^6^ RD cells maintained under growth condition were crosslinked followed by purification of nuclei, and chromatin shearing by sonication (eight times, 20 s each). Sheared chromatin was incubated overnight with antibodies against sXBP1 (Cell Signaling Technology) or Histone H3 (Cell Signaling Technology) followed by incubation with Protein G magnetic beads. Normal Rabbit IgG (Cell Signaling Technology) was used as a negative control for the immunoprecipitation experiment. Magnetic beads were briefly washed, and chromatin DNA was eluted, reverse crosslinked, and purified. Purified DNA was analyzed for enrichment of 100-150 bp sequences by quantitative real time-PCR (40 cycles) or semi-quantitative standard PCR using primer set (Forward: AGC GGT TAA CAA ACT GTT CAG A, Reverse: CCA GAA TAC GAT GTT CCT GGG T) designed for binding sites in the BMPR1A enhancer region. The fold change between negative control IgG and anti-sXBP1 groups was calculated using 2−ΔΔCt formula normalized by signal from 2% input.

### RNA-sequencing and data analyses

Total RNA from control, IRE1α, or XBP1 knockdown RD cells was extracted using TRIzol reagent (Thermo Fisher Scientific) using RNeasy Mini Kit (QIAGEN) according to the manufacturers’ protocols. The mRNA-seq library was prepared using poly (A)-tailed enriched mRNA at the UT Cancer Genomics Center using the KAPA mRNA HyperPrep Kit protocol (KK8581, Roche, Holding AG, Switzerland) and KAPA Unique Dual-indexed Adapter kit (KK8727, Roche). The Illumina NextSeq550 was used to produce 75 base paired-end mRNA-seq data at an average read depth of ∼38 M reads/sample. RNA-seq fastq data were processed using CLC Genomics Workbench 20 (Qiagen). Illumina sequencing adapters were trimmed, and reads were aligned to the mouse reference genome Homo Sapiens (GRCh38/hg38). Normalization of RNA-seq dataset was performed using a trimmed mean of M values. Genes with |Log_2_FC| ≥ 0.25 and p-value < 0.05 were assigned as differentially expressed genes (DEGs) and represented in volcano plot using ggplot function in R software (v 4.2.2). Pathway enrichment analysis associated with the up-regulated and down-regulated genes were identified using the Metascape gene annotation and analysis tool (metascape.org) as described ^69,70^. Heatmaps were generated using the heatmap.2 function ^71^ using z-scores calculated based on transcripts per million (FPKM) values. TPM values were converted to log (TPM□□+□□1) to handle zero values. Genes involved in specific pathways were manually selected for heatmap expression plots. The average absolute TPM values for the control group are provided in **Supplementary Table S2**. All the raw data files can be found on the NCBI SRA repository using the accession code PRJNA1435360.

### Xenograft growth mouse model

Six-week-old male Nu/Nu mice were purchased from Charles River Laboratories. RD cells expressing Tet-On SCR or XBP1 shRNA along with luciferase were suspended in Matrigel (Corning) and approximately 5 × 10^6^ cells in 50% Matrigel (Sigma-Aldrich) were injected subcutaneously into the flank of each mouse. Body weight and tumor growth were monitored weekly, and tumor length (L) and width (W) were measured. Tumor volume was calculated using the formula 0.5 × L × W^2. Tumor growth was further assessed by bioluminescence imaging using a Xenogen IVIS® system (PerkinElmer). When tumor volumes reached approximately 50-100 mm³, mice were randomly assigned to 2 groups. One group received standard chow (control), and the other group received doxycycline-containing chow. Mice were euthanized after 4 weeks of doxycycline treatment. For pharmacological studies, approximately 5 × 10^6^ RD cells suspended in 50% Matrigel were injected subcutaneously into the flank of each mouse. When the tumor volume reached approximately 50-100 mm³, mice received intraperitoneal injections of 4μ8C (10 mg/kg; diluted in 5% DMSO and 95% corn oil) or vehicle (control) every other day. Mice were euthanized after 4 weeks of vehicle or 4μ8C treatment. The animal protocol (PROTO201900043) was approved by the Institutional Animal Care and Use Committee (IACUC) of the University of Houston. We have complied with all relevant ethical regulations for animal use.

### Histology and Immunohistochemistry

Tumors were isolated and processed for histological and biochemical analyses. Tumor samples were fixed in 4% paraformaldehyde overnight at 4°C, washed with cold PBS, and embedded in Frozen Section Compound (FSC22, Leica, USA). Cryosections of 10 µm thickness were prepared and stained with hematoxylin and eosin (H&E). Sections were also immunostained for Ki67 and myogenin protein. Nuclei were counterstained by DAPI.

### Statistics and Reproducibility

Sample size was determined by a priori power analysis based on estimates of the standard deviation (s.d.) and effect size obtained from prior experiments using the same procedures. For animal studies, sample size was estimated based on the anticipated availability of nude mice. Power calculations indicated a minimal sample size of eight animals per group. Considering a potential attrition rate of approximately 10%, six mice were initially assigned to each group. For some experiments, three to four animals per group were sufficient to detect statistically significant differences. Animals of the same sex and age were used to minimize physiological variability and reduce variation around the mean. Animals were randomly allocated to experimental groups, and animals from different cages were included to avoid cage-specific bias. Investigators were blinded to group allocation during experiments and outcome assessment through the use of coded identifiers until completion of the final data analysis.

Exclusion criteria were predefined in consultation with the IACUC and based on experimental endpoints. Animals were excluded from analysis in cases of death, skin injury, illness, or body weight loss exceeding 10%. Tumor samples were excluded if technical artifacts occurred, such as freezing artifacts in histological sections or failure to obtain protein of sufficient quality or quantity for downstream analyses. No statistical methods were used to predetermine sample size beyond power analysis, but the sample sizes are consistent with those commonly used in the field. Data distribution was assumed to be normal, but this was not formally tested. Statistical analyses were performed as indicated in the corresponding figure legends. Data are presented as mean ± s.d. Comparisons between two groups were performed using an unpaired two-tailed Student’s *t*-test. A *p* value < 0.05 was considered statistically significant.

## Data availability statement

Raw data files for RNA-seq experiment can be found on the NCBI SRA repository using the accession code PRJNA1435360. All other data are included with this manuscript and also available from the corresponding author on reasonable request.

## Acknowledgements

We are grateful to Dr. Peter J Houghton of The University of Texas Health Science Center at San Antonio, Texas for providing RH36 and RH41 cell lines. We thank technical support from the Cancer Prevention and Research Institute of Texas (CPRIT RP180734) core for RNA-seq experiments.

## Authors’ contributions

AK, BG, and MVT designed the work. ATV, ASJ, PTH, and MTS performed the experiments and analyzed the results. ATV, ASJ, and AK wrote the manuscript. All authors edited and finalized the manuscript.

