## Supplementary material for "Targeting the IRE1α-XBP1 signaling axis impairs tumor growth and promotes myogenic differentiation in rhabdomyosarcoma": Figures S1-S7; Tables S1 and S2

This file contains **Figures S1-S7** and **Tables S1 and S2.**

### Supplemental Figures and Legends

#### FIGURE S1

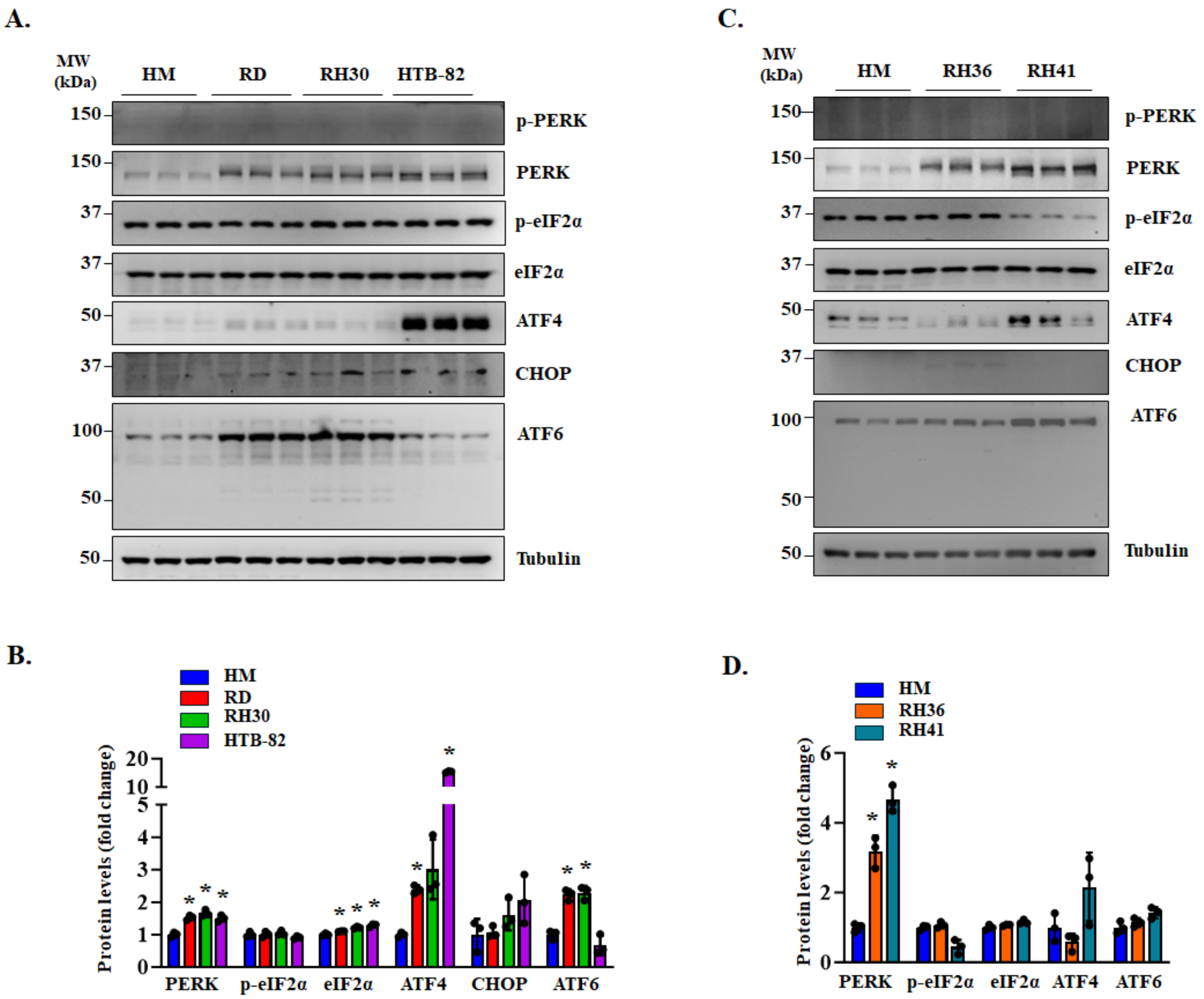


**FIGURE S1. Expression of ER stress/UPR markers in RMS cell lines. (A, C)** Immunoblots and **(B, D)** densitometry analysis showing the levels of phosphorylated PERK (p-PERK), total PERK, p-eIF2α, total eIF2α, ATF4, CHOP, ATF6 and Tubulin protein in human myoblasts (HM), and RD, RH30, HTB-82, RH36, and RH41 cell lines. n=3 biological replicates per group. Data are presented as mean ± SD. *$p<0.05$, values significantly different from HM analyzed by unpaired Student *t* test.

FIGURE S2

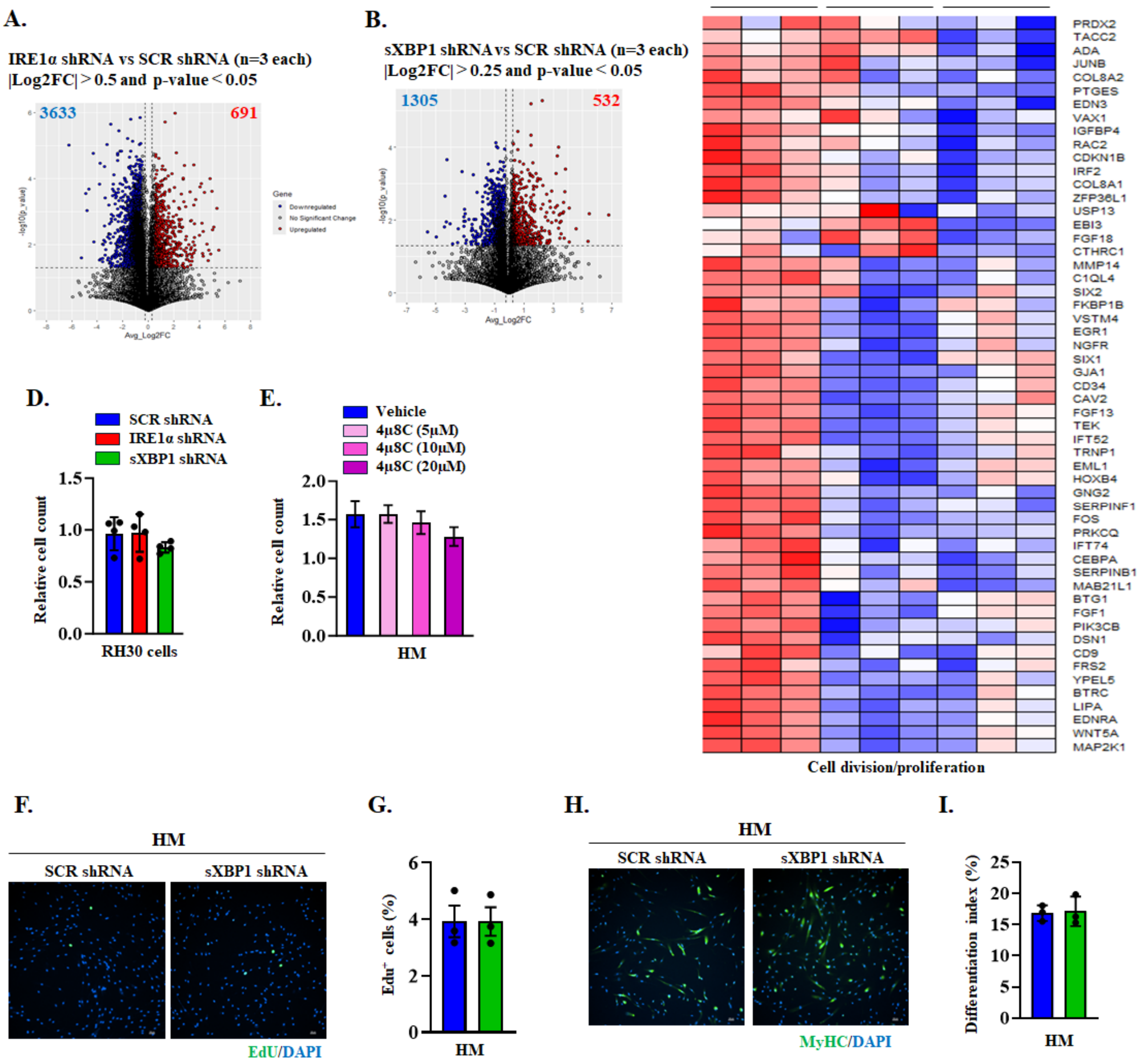


**FIGURE S2. Effect of knockdown of IRE1α or sXBP1 on proliferation of RMS cells.** Volcano plots showing differentially expressed genes in **(A)** IRE1α shRNA and **(B)** sXBP1 shRNA expressing RD cells compared with SCR shRNA expressing RD cultures. **(C)** Heatmap generated after analysis of RNA-seq dataset show relative mRNA levels of various molecules involved in cell division and proliferation-related genes in RD cultures expressing scrambled (SCR), IRE1α, or sXBP1 shRNA. **(D)** Relative number of RH30 cells expressing SCR, IRE1α, or sXBP1 shRNA after culturing for 72 h in growth medium. **(E)** Relative number of human myoblasts (HM) after treatment with indicated concentrations of 4μ8C for 4 days in growth medium. **(F)** Representative images of control and sXBP1 knockdown HM after performing EdU staining. Scale bar, 100 μm. **(G)** Quantification of percentage of $EdU^{+}$ cells in control and sXBP1 knockdown HM cultures. **(H)** Representative images of control and sXBP1 knockdown HM after performing immunostaining for MyHC. Scale bar, 100 μm. **(I)** Quantification of differentiation index in control and sXBP1 knockdown HM cultures. n=3 biological replicates per group. Data are presented as mean ± SD. No significant difference was observed using unpaired Student t test.

#### FIGURE S3

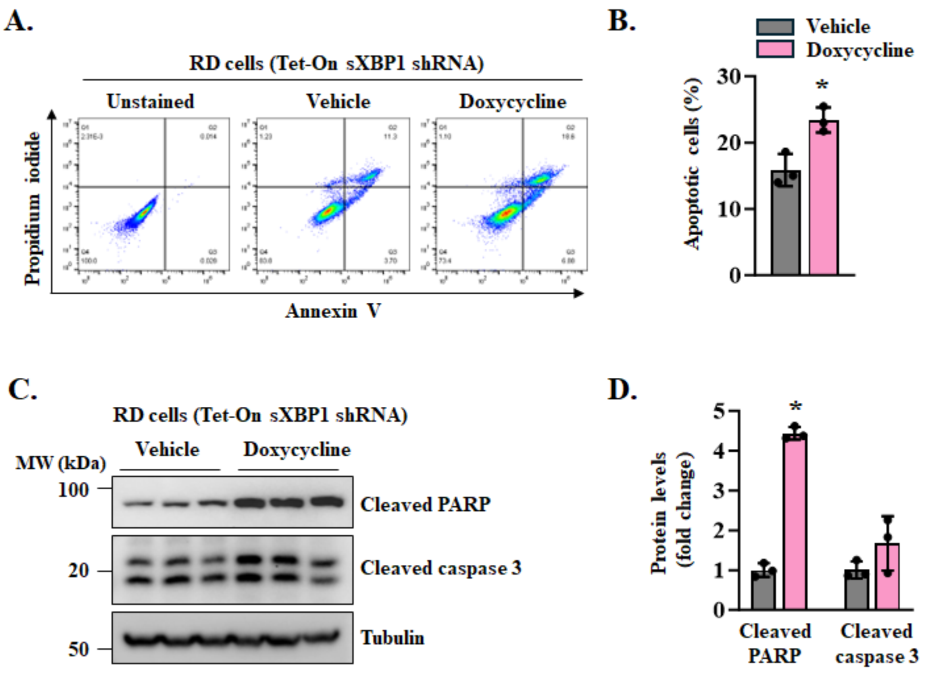


**FIGURE S3. Inducible knockdown of XBP1 promotes apoptosis in RD cells.** RD cells stably expressing Tet-On XBP1 shRNA were treated with vehicle alone or 1μg/ml doxycycline for 5days followed by propidium iodide (PI) and Annexin V staining and FACS analysis. **(A)** Scatter plots and **(B)** quantification of proportion of apoptotic cells in control and doxycycline-treated RD cultures. n=3 biological replicates per group. Data are presented as mean ± SD. *p<0.05, values significantly different from vehicle-treated control cultures analyzed by unpaired Student *t* test. **(C)** Immunoblots and **(D)** densitometry analysis showing levels of cleaved PARP and cleaved Caspase-3 protein in control and doxycycline-treated RD cultures. n=3 biological replicates per group. Data are presented as mean ± SD. *p<0.05, values significantly different from vehicle-treated control cultures analyzed by unpaired Student *t* test.

#### FIGURE S4

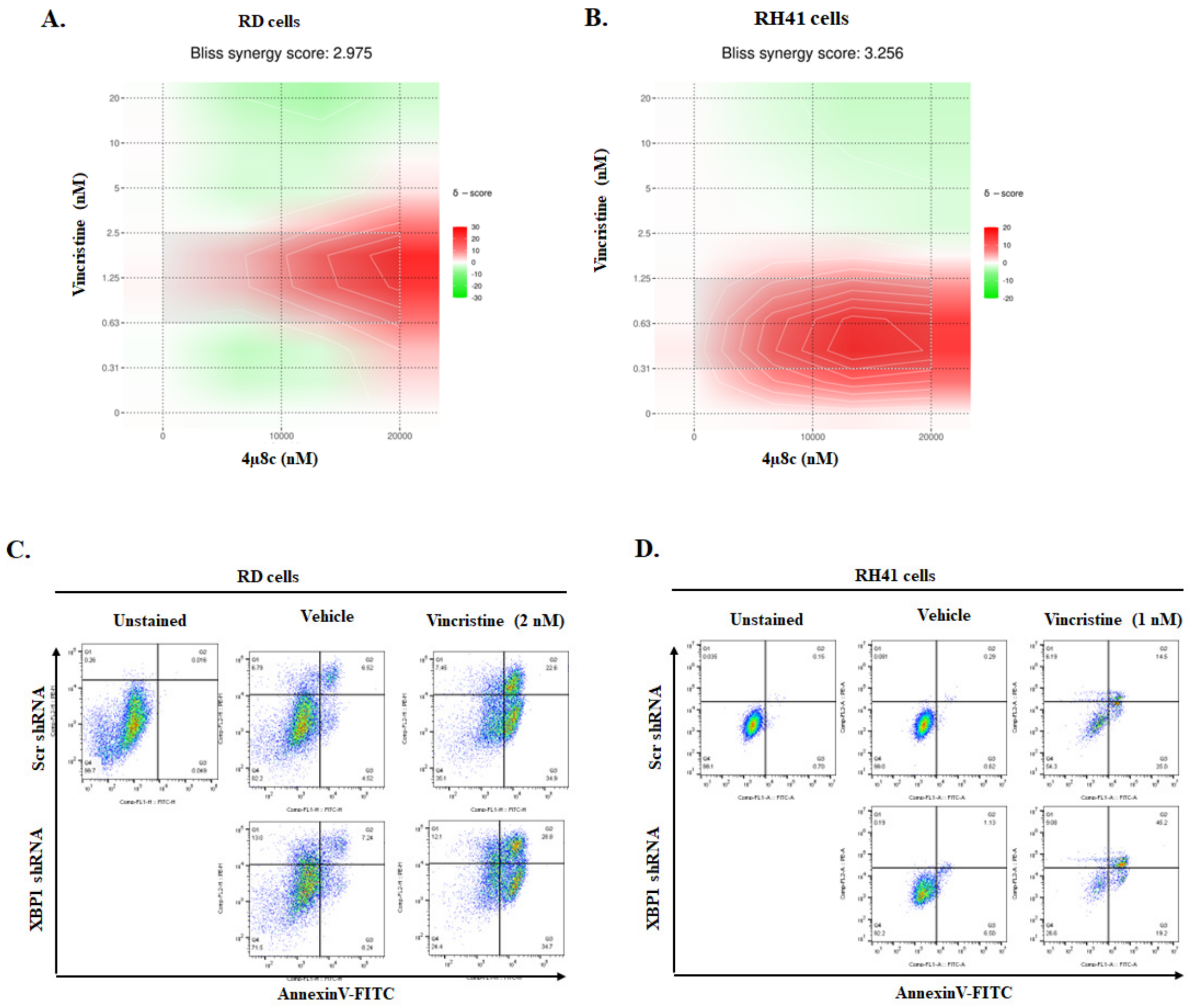


**FIGURE S4. Inhibition of XBP1 increases vincristine-induced apoptosis in RMS cells.** Vincristine and 4μ8C interaction analysis was performed using the Bliss independence model, and synergy scores were calculated using SynergyFinder. Drug combination matrix showing the effects of varying concentrations of vincristine and 4μ8C on the viability of (A) RD cells, **(B)** RH41 cells. Positive values of Bliss scores suggest synergistic interactions between vincristine and 4μ8C. Scatter plots of FACS analysis presented here demonstrate that knockdown of XBP1 increases the vincristine-induced apoptosis in **(C)** RD cells and **(D)** RH41 cells. Quantification is provided in main Figure 4F.

**FIGURE S5**

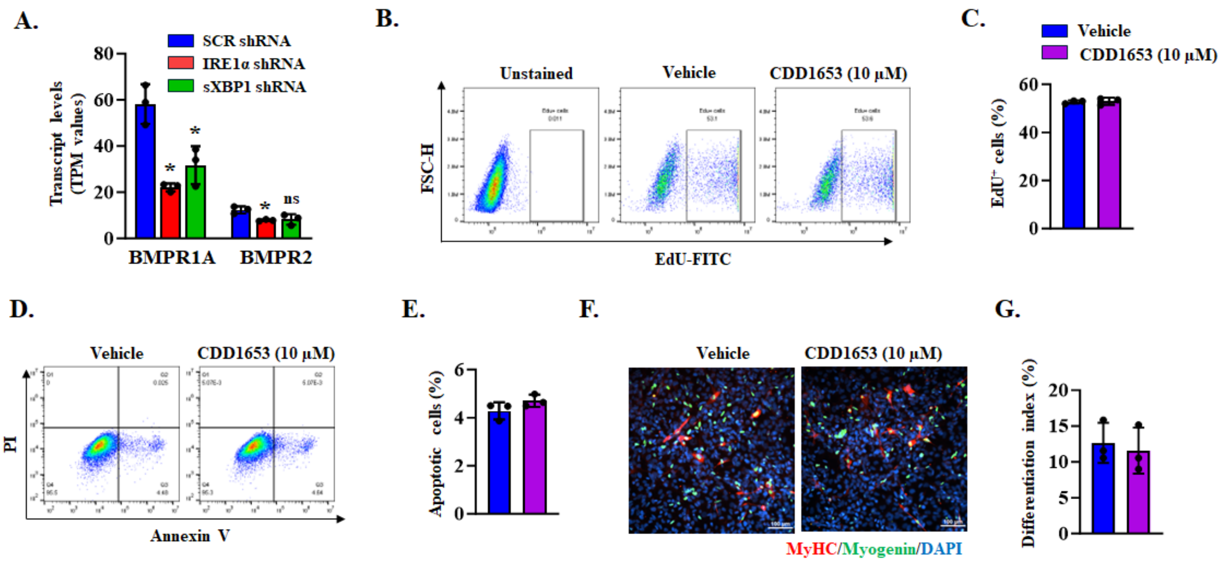


**FIGURE 5. Effect of BMPR2 inhibition on RD cells. (A)** TPM values from RNA-seq dataset for the expression of BMPR1A and BMPR2 in RD cells expressing scrambled (SCR), IRE1α, or sXBP1 shRNA. n=3 biological replicates per group. Data are presented as mean ± SD. *$p < 0.05$, values significant different from corresponding SCR shRNA expressing RD cultures. **(B)** Scatter plots of flow cytometry, and **(C)** quantification of $EdU^+$ cells in cultures treated with vehicle alone or 10µM CDD1653. **(D)** Scatter plots, and **(E)** quantification of Annexin $V^+$ cells in cultures treated with vehicle alone or 10µM CDD1653. **(F)** Representative images of vehicle and CDD1653-treated RD culture after immunostaining for myogenin and MyHC proteins. Nuclei were stained with DAPI. **(G)** Quantification of differentiation of index in control and CD1653-treated RD cultures. n=3 biological replicates per group. Data are presented as mean ± SD. No significant difference was observed using unpaired Student *t* test.

FIGURE S6

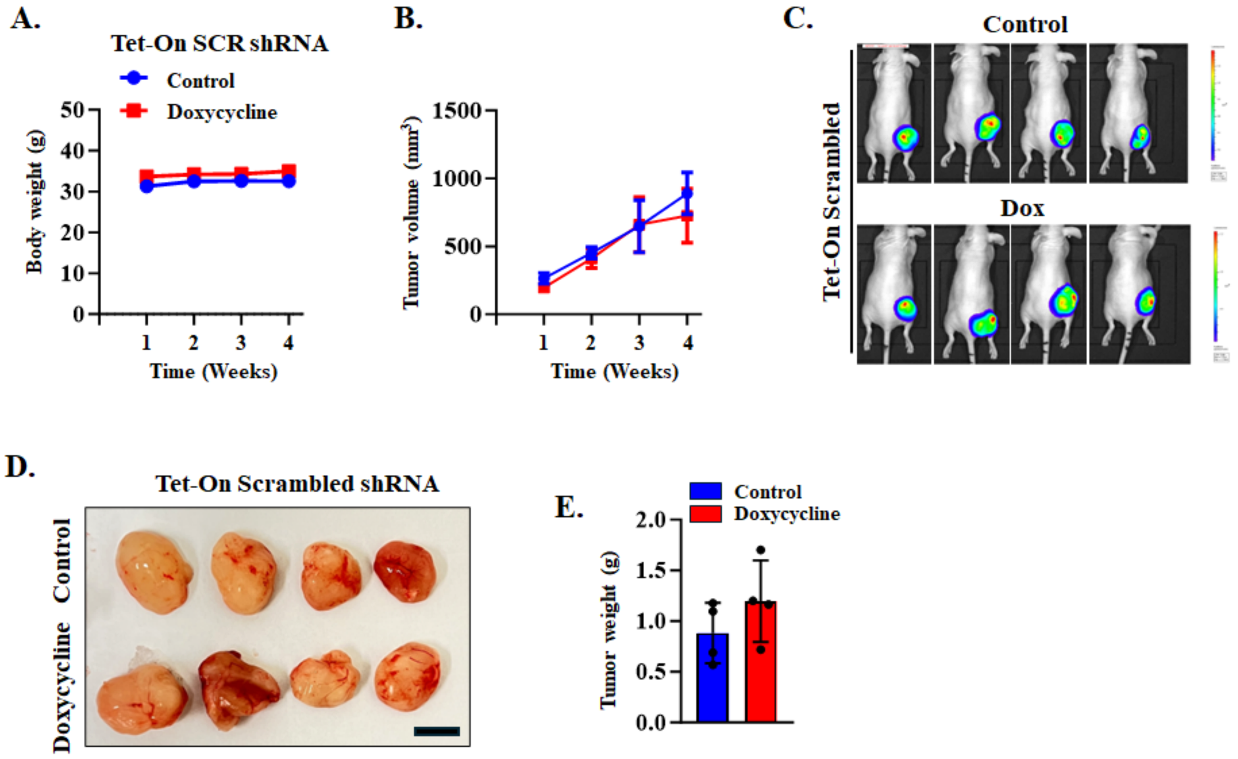


**FIGURE S6. Treatment with doxycycline alone does not alter tumor growth in RD xenografts. (A)** Body weight of mice inoculated with Tet-On scrambled (SCR) shRNA expressing RD cells fed with normal chow or doxycycline-containing chow. **(B)** Average tumor volume in mice fed with normal chow or doxycycline containing chow. n=4 mice in each group. Data are presented as mean ± SD. No significant difference was observed using unpaired Student *t* test. **(C)** Bioluminescence imaging showing presence RD xenograft in control and doxycycline-treated nude mice. **(D)** Images of tumors at the time of euthanizing the mice between control and doxycycline-treated groups. Scale bar, 1cm. **(E)** Quantification of wet weight of tumors in control and doxycycline-treated mice. n=4 mice per group. Data are presented as mean ± SD. No significant difference was observed using unpaired Student *t* test.

#### FIGURE S7

##### Figure 1D

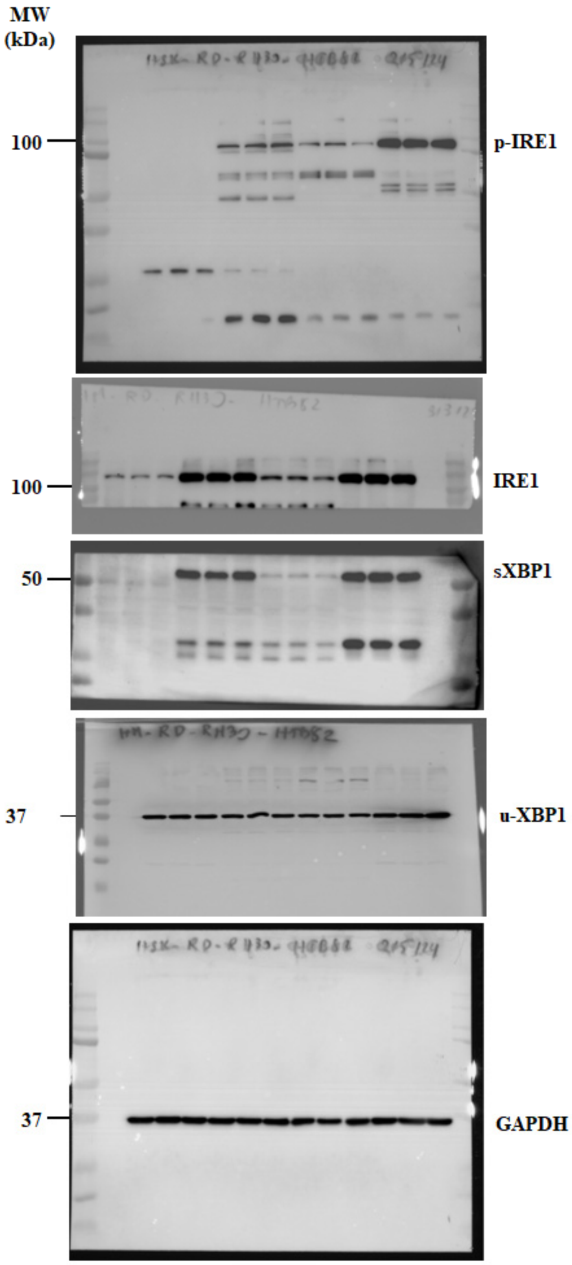


##### Figure 1F

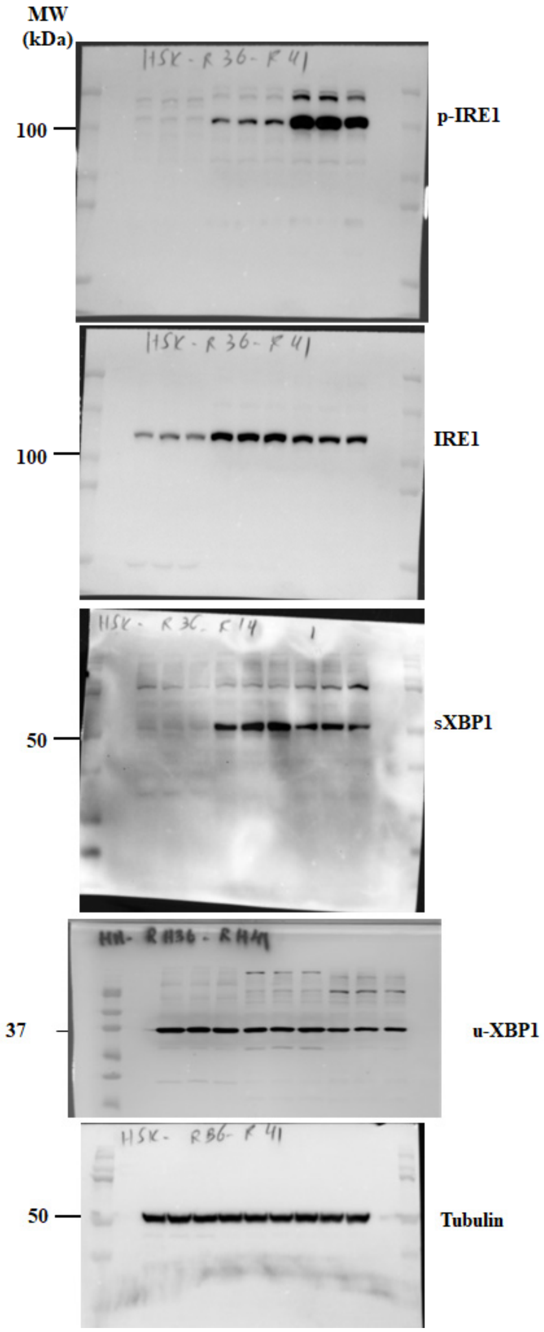

#### FIGURE S7 (continuation)

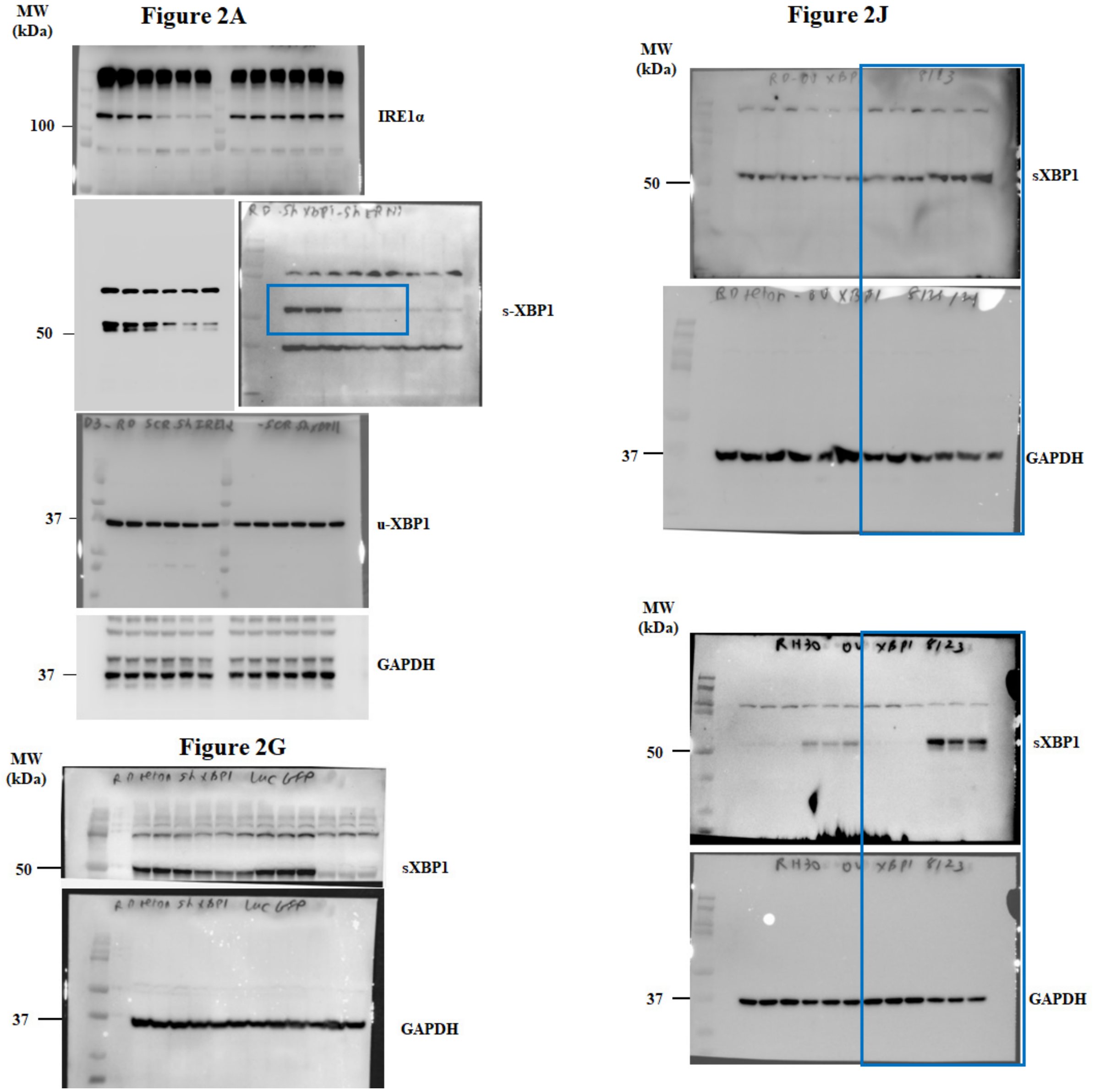

**FIGURE S7 (continuation)**

**Figure 3C**

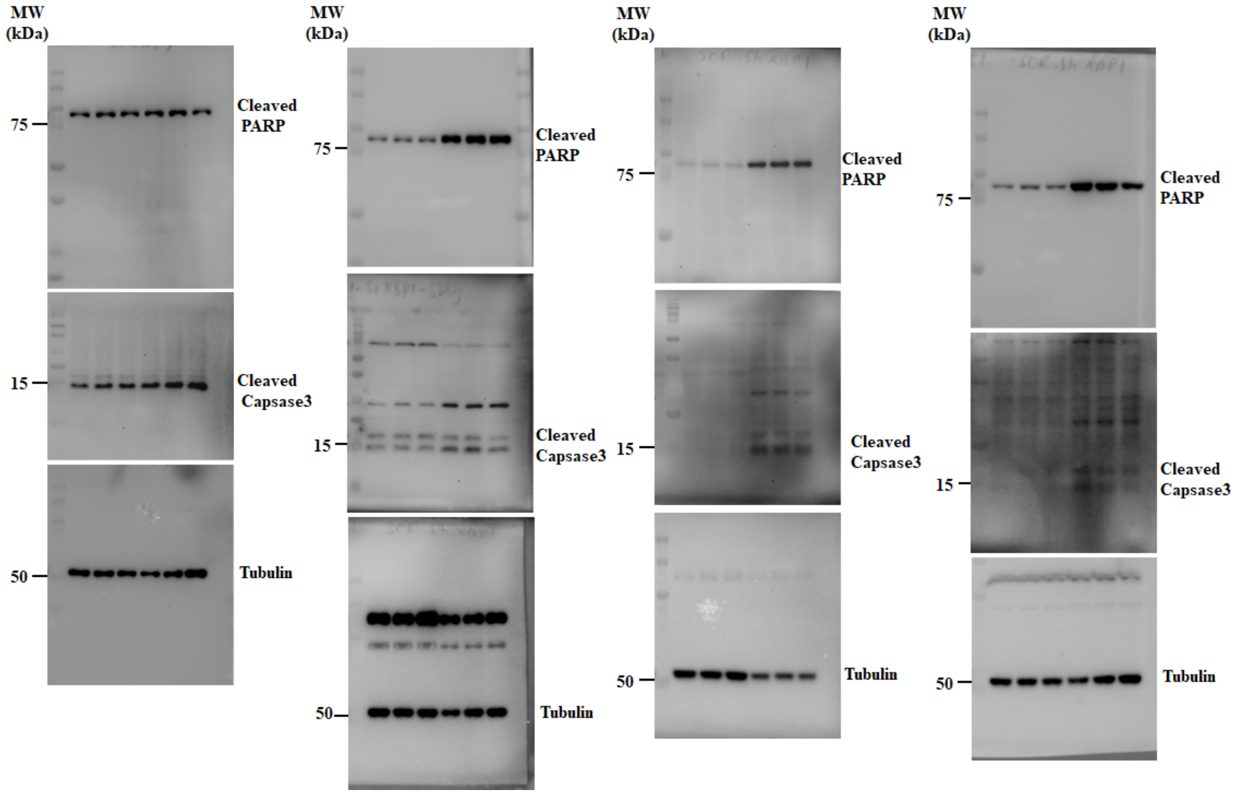

**FIGURE S7 (continuation)**

#### Figure 3G

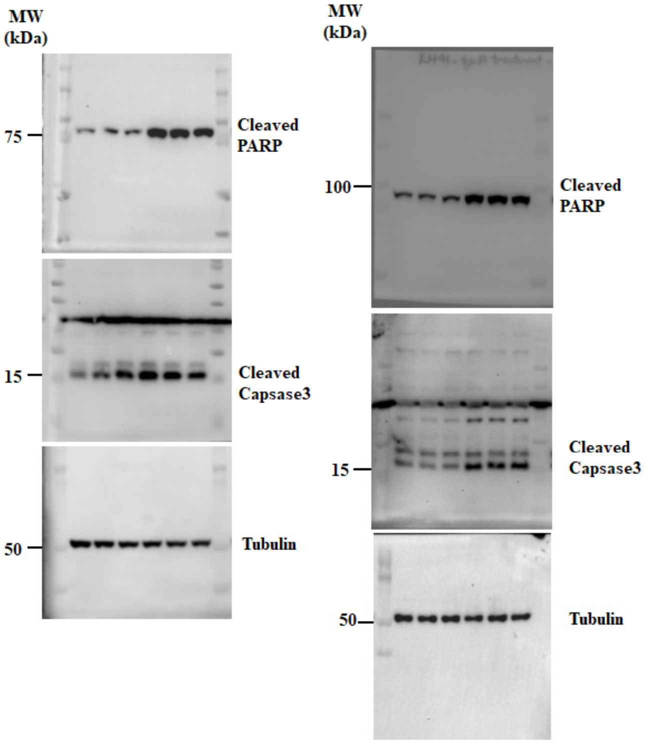


#### Figure 5B

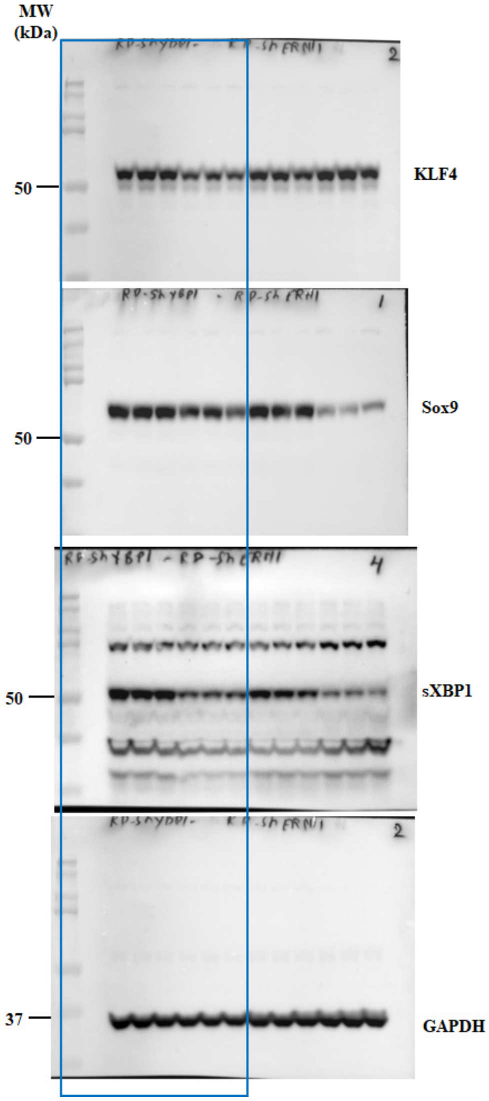

#### FIGURE S7 (continuation)

##### Figure 6L

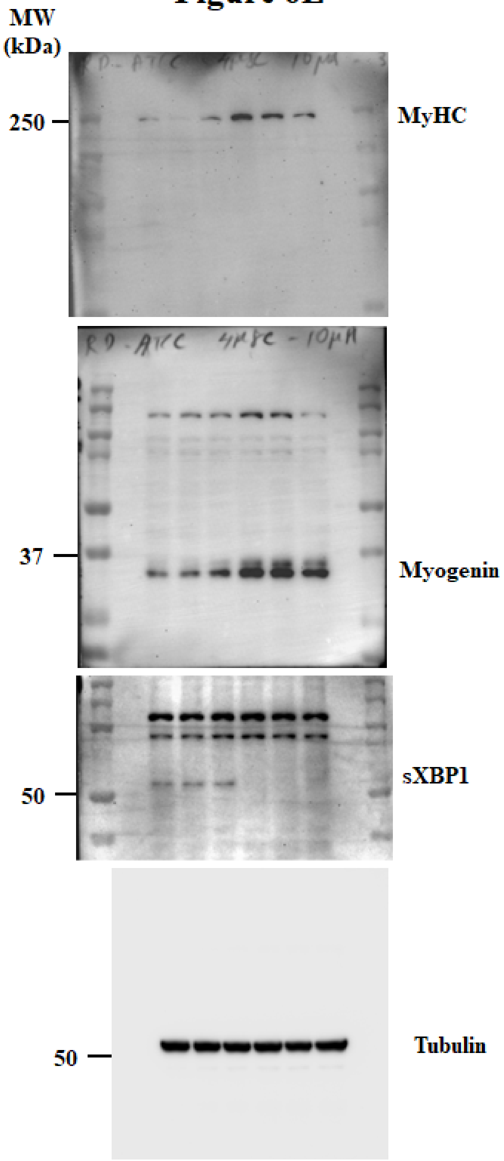

**FIGURE S7 (continuation)**

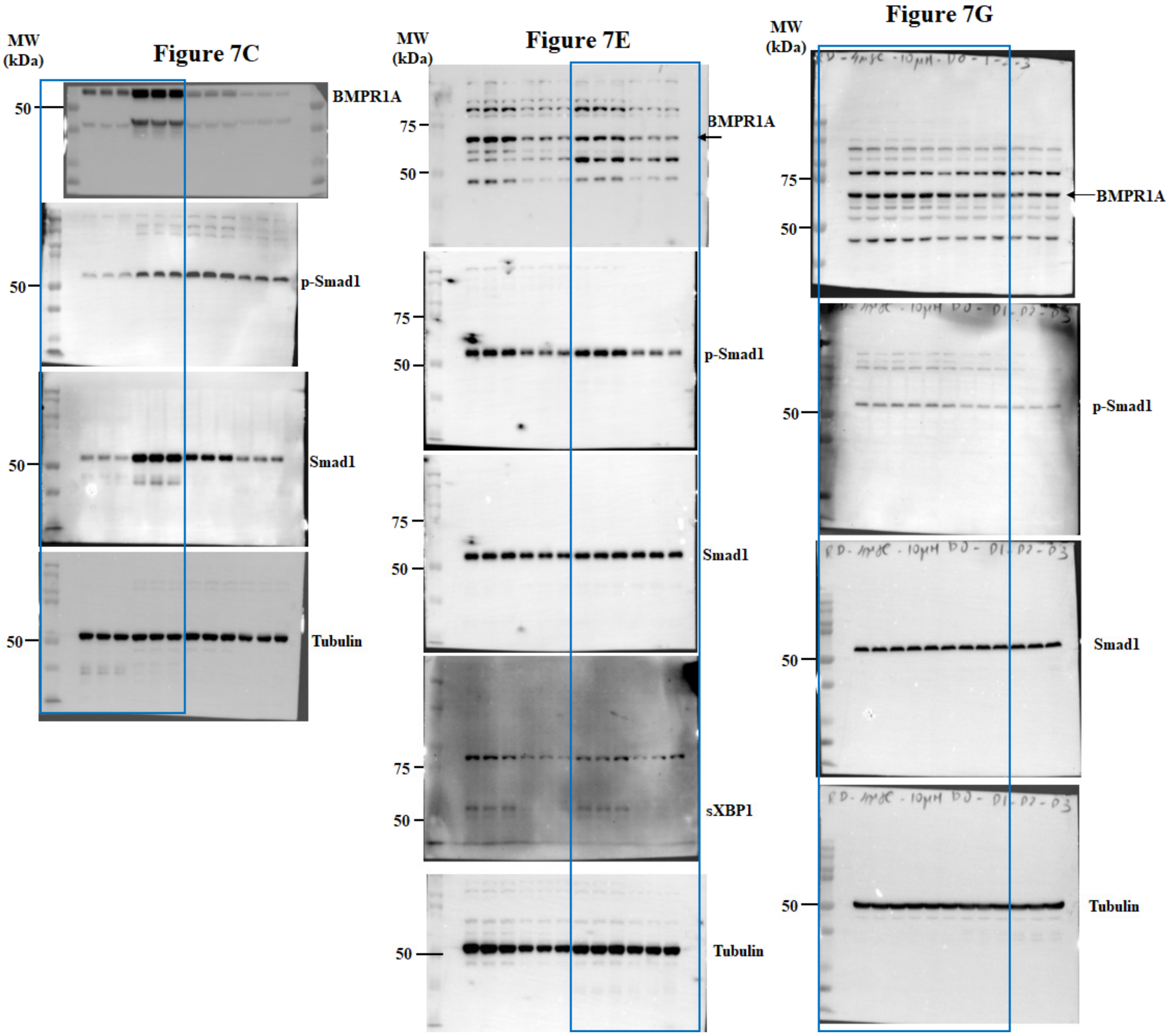

**FIGURE S7 (continuation)**

**Figure 7I**

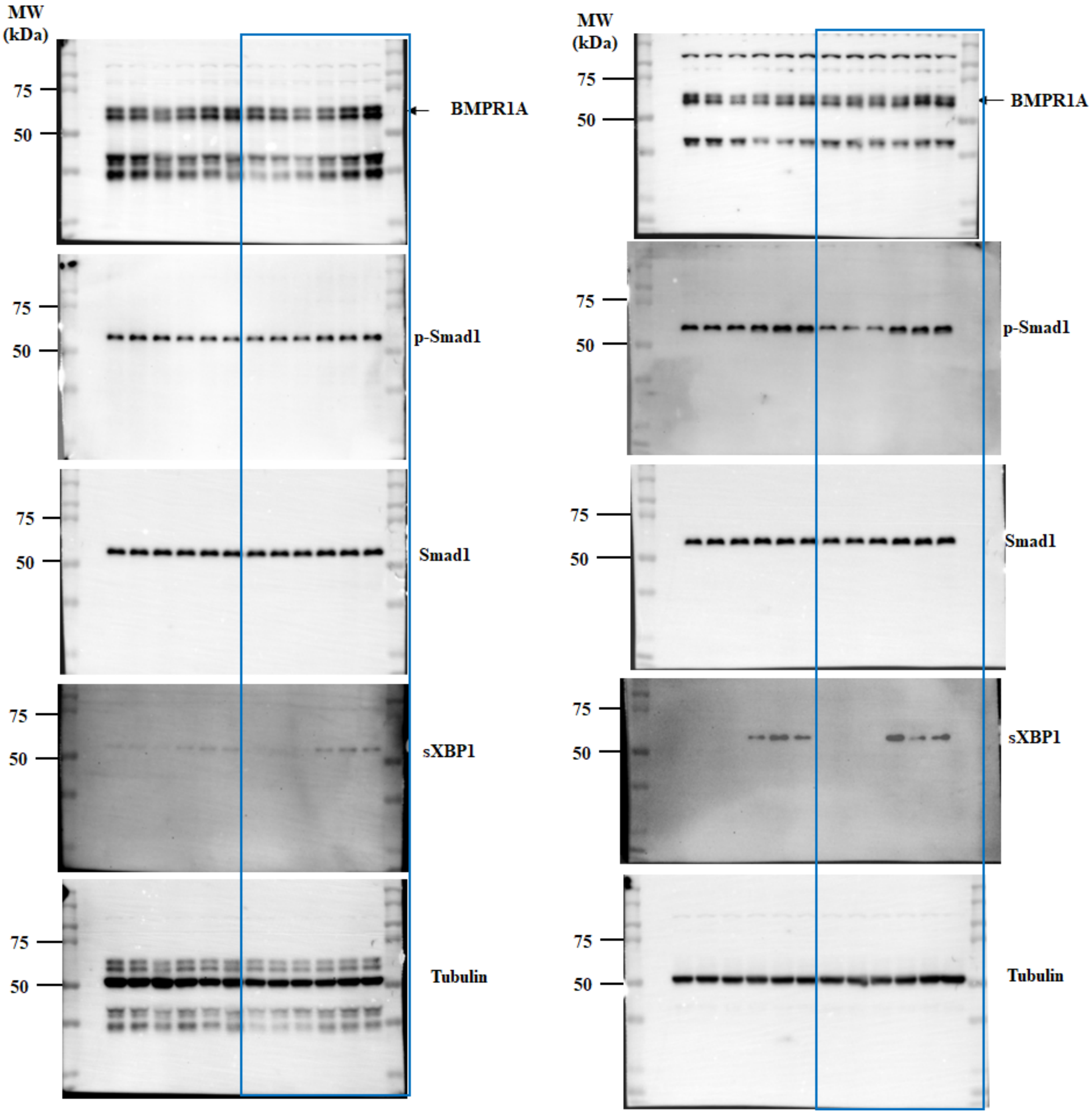

**FIGURE S7 (continuation)**

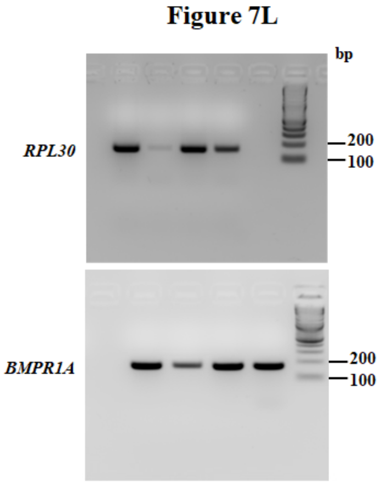


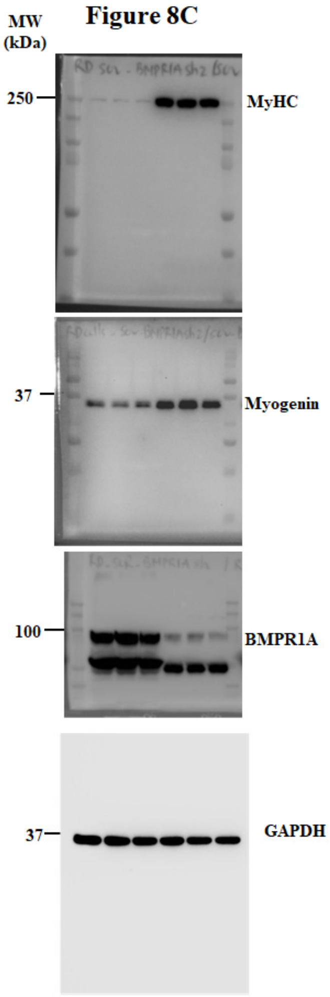


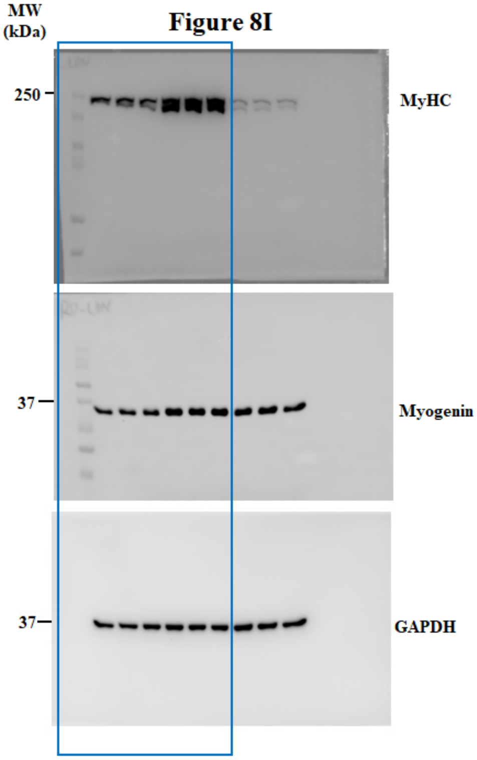

### FIGURE S7 (continuation)

#### Figure 9K

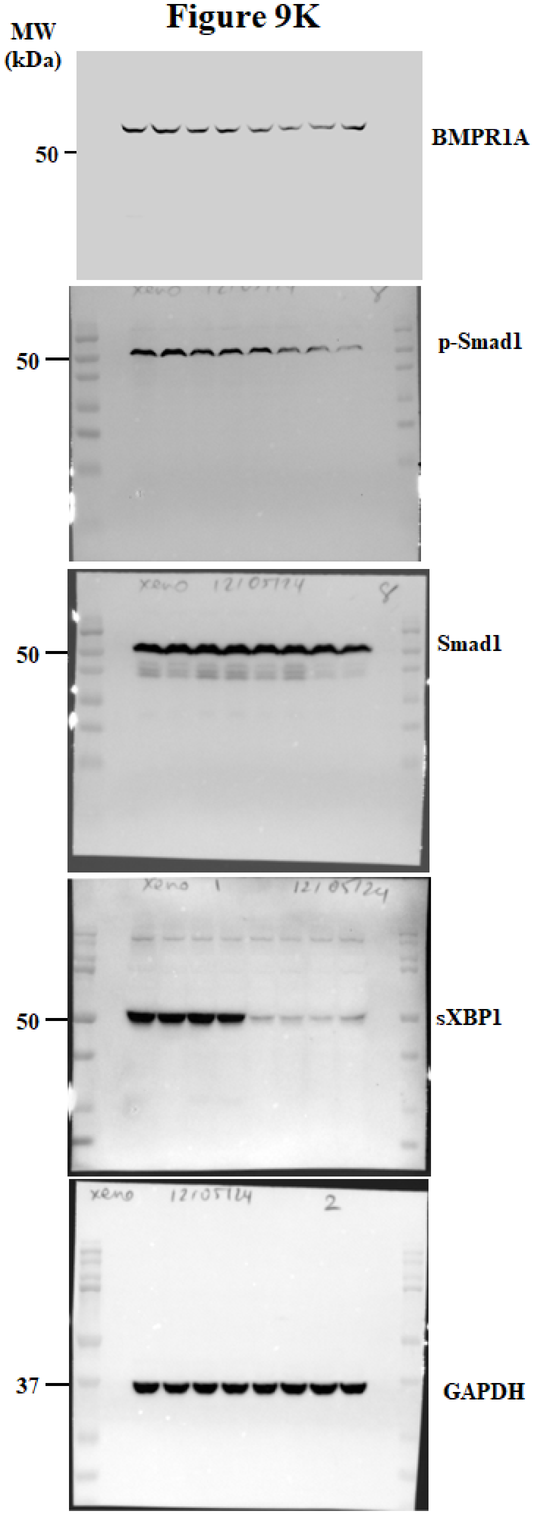


#### Figure S1A

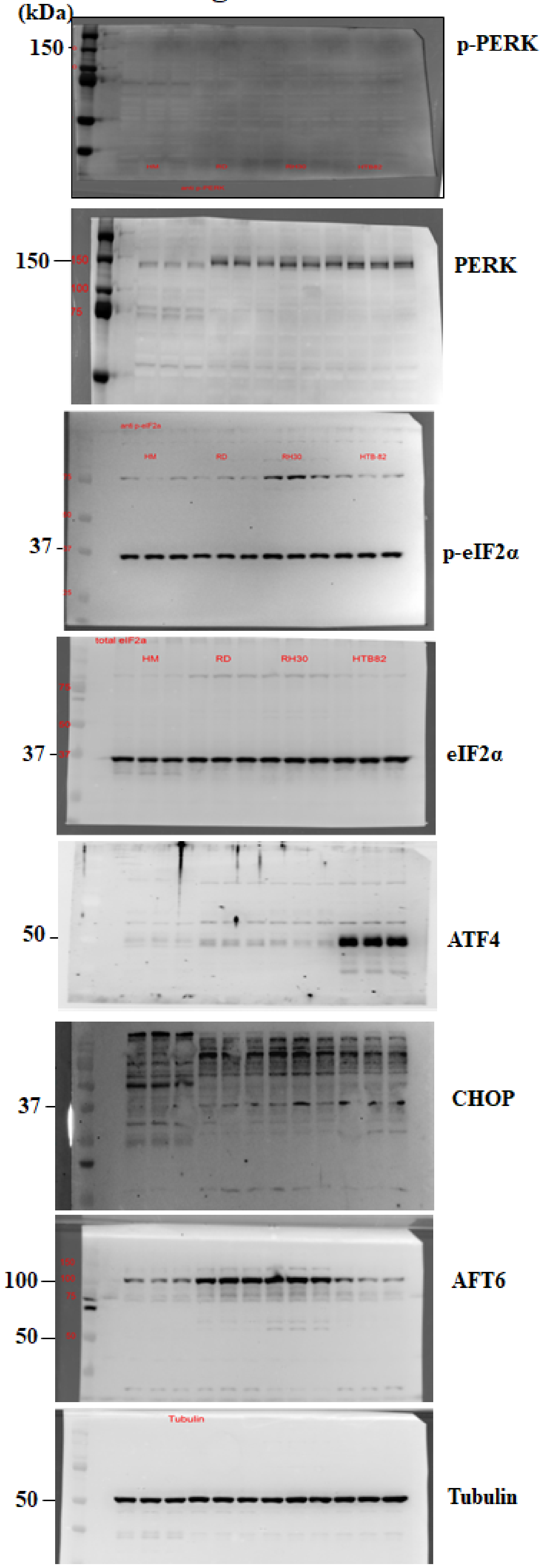

**FIGURE S7 (continuation)**

**Figure S1C**

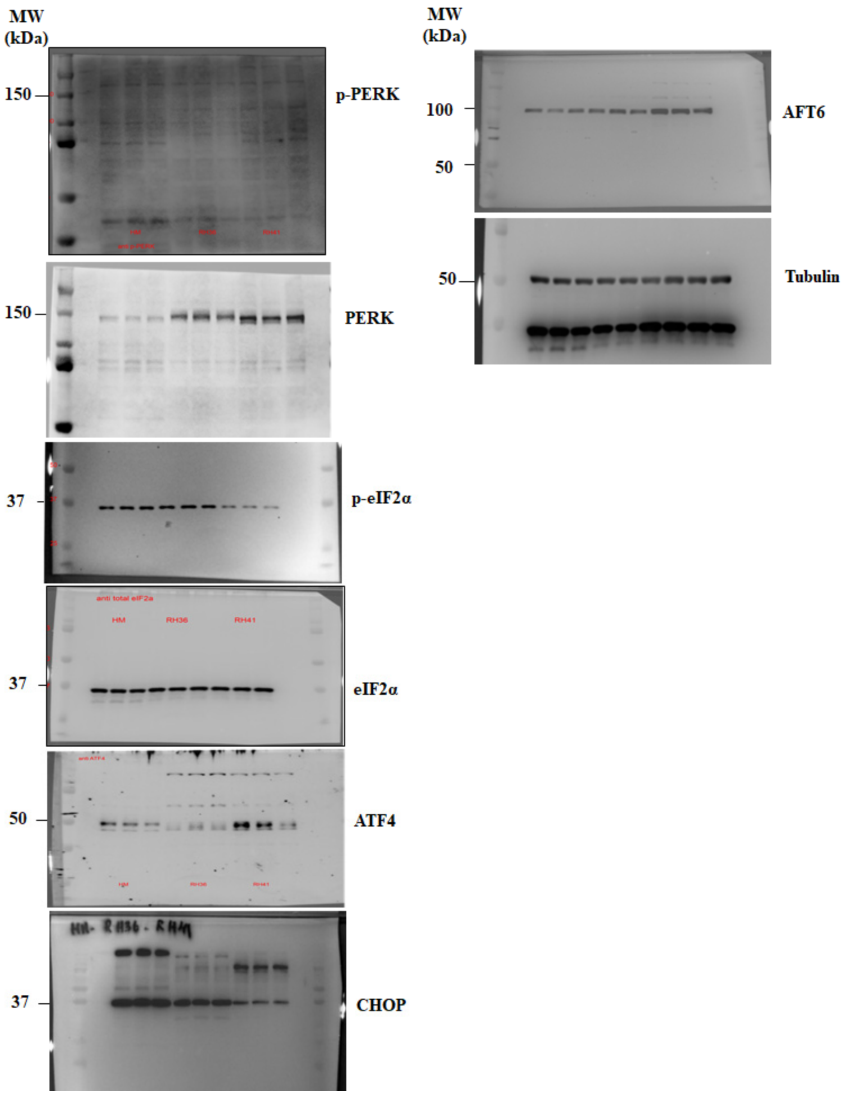


**Figure S3C**

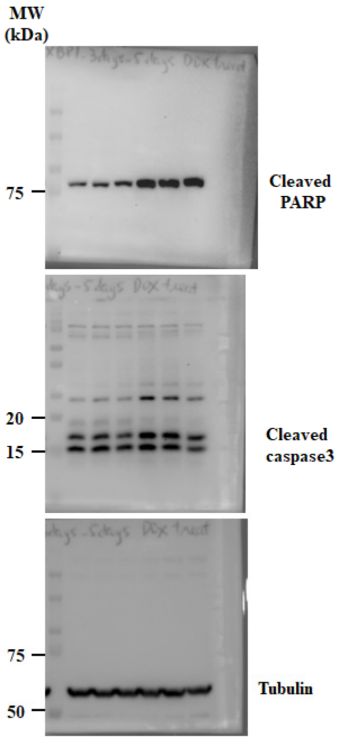


**FIGURE S7. Uncropped gel images**. Original uncropped images of western blot and agarose gels.

**Table S1. List of antibodies, source, and dilution used.** ChIP, chromatin immunoprecipitation; IF, immunofluorescence; WB, western blot.

| Antibody | Dilution | Source | Identifier |
|---|---|---|---|
| Rabbit-anti-p-ERN1 | 1:1000 (WB) | Abnova | # PAB12435 |
| Rabbit-anti-IRE1 alpha | 1:1000 (WB) | Cell Signaling Technology | # 3294 |
| Rabbit-anti-sXBP1 | 1:1000 (WB)/1:50 (ChIP) | Cell Signaling Technology | # 40435 |
| Rabbit-anti-alpha Tubulin | 1:1000 (WB) | Cell Signaling Technology | # 2144 |
| Rabbit-anti-GAPDH | 1:1000 (WB) | Cell Signaling Technology | # 2118 |
| Rabbit-anti-Cleaved PARP | 1:1000 (WB) | Cell Signaling Technology | # 5625 |
| Rabbit-anti-Cleaved Caspase 3 | 1:1000 (WB) | Cell Signaling Technology | # 9661 |
| Rabbit-anti-KLF4 | 1:1000 (WB) | Cell Signaling Technology | # 12173 |
| Rabbit-anti-Sox9 | 1:1000 (WB) | Cell Signaling Technology | # 82630 |
| Mouse-anti-Myosin heavy chain | 1:500 (WB)/1:50 (IF) | DSHB | # MF20 |
| Mouse-anti-Myogenin | 1:500 (WB)/1:50 (IF) | Invitrogen | # MA5-11486 |
| Rabbit-anti-BMPR1A | 1:1000 (WB) | Proteintech | #82928-1-RR |
| Rabbit-anti-p-Smad1 | 1:1000 (WB) | Cell Signaling Technology | # 5753 |
| Rabbit-anti-Smad1 | 1:1000 (WB) | Cell Signaling Technology | # 6944 |
| Rabbit-anti-Ki67 | 1:500 (IF) | Invitrogen | #MA5-14520 |
| Rabbit-anti-XBP1 (For uXBP1) | 1:1000 (WB) | Abcam | # ab37152 |
| Rabbit-anti-p-PERK | 1:1000 (WB) | Cell Signaling Technology | # 3179S |
| Rabbit-anti-PERK | 1:1000 (WB) | Cell Signaling Technology | # 3192 |
| Mouse-anti-CHOP | 1:1000 (WB) | Cell Signaling Technology | # 2895S |
| Rabbit-anti-p-eIF2a | 1:1000 (WB) | Cell Signaling Technology | # 9721 |
| Rabbit-anti-eIF2a | 1:1000 (WB) | Cell Signaling Technology | # 9722 |
| Rabbit-anti-ATF4 | 1:1000 (WB) | Cell Signaling Technology | # 11815 |
| Rabbit-anti-ATF6 | 1:1000 (WB) | Cell Signaling Technology | # 65880 |
| Anti-rabbit Ig, HRP-linked | 1:2000 (WB) | Cell Signaling Technology | # 7074S |
| Anti-mouse Ig, HRP-linked | 1:2000 (WB) | Cell Signaling Technology | # 7076 |
| Anti-Mouse IgG2 AF594 | 1:1000 (IF) | Invitrogen | # A211135 |
| Anti-Mouse IgG2b AF488 | 1:1000 (IF) | Invitrogen | # A21141 |
| Anti-Rabbit IgG AF488 | 1:1000 (IF) | Invitrogen | # A32731 |

**Table S2. Average TPM scores in control RD cultures expressing SCR shRNA.**

| Gene | Avg TPM value | Gene | Avg TPM value | Gene | Avg TPM value | Gene | Avg TPM value |
|---|---|---|---|---|---|---|---|
| KLF4 | 6.019 | CAV3 | 0.111 | PTGES | 4.079 | IFT52 | 47.469 |
| KLF7 | 4.842 | MYH1 | 0.185 | EDN3 | 192.454 | TRNP1 | 5.090 |
| LGR5 | 18.897 | MYH4 | 0.286 | VAX1 | 26.690 | EML1 | 18.746 |
| ALDH1A3 | 4.151 | MYH2 | 0.117 | IGFBP4 | 5.031 | HOXB4 | 15.241 |
| SOX7 | 4.034 | MYOG | 11.005 | RAC2 | 4.628 | GNG2 | 88.089 |
| ZEB1 | 11.188 | MYL9 | 74.677 | CDKN1B | 18.663 | SERPINF1 | 58.095 |
| BMI1 | 17.633 | MKX | 0.009 | IRF2 | 17.648 | FOS | 2.901 |
| NGFR | 324.698 | NPNT | 0.704 | COL8A1 | 10.102 | PRKCQ | 1.741 |
| CD34 | 7.666 | MYL6 | 1301.628 | ZFP36L1 | 54.502 | IFT74 | 8.142 |
| SOX9 | 13.691 | MEF2B | 0.605 | USP13 | 12.948 | CEBPA | 3.303 |
| ITGA7 | 13.956 | MYL1 | 8.193 | EBI3 | 0.176 | SERPINB1 | 10.740 |
| MYL4 | 0.508 | ACTC1 | 58.935 | FGF18 | 0.943 | MAB21L1 | 1.678 |
| TNNC1 | 0.464 | BMPR1A | 58.138 | CTHRC1 | 208.887 | BTG1 | 42.452 |
| LAMA1 | 0.021 | BMPR2 | 12.381 | MMP14 | 52.950 | FGF1 | 2.081 |
| TTN | 0.053 | SMAD1 | 40.768 | C1QL4 | 1.618 | PIK3CB | 4.232 |
| MYOZ2 | 0.057 | SMAD2 | 41.776 | SIX2 | 90.733 | DSN1 | 38.102 |
| ANKRD1 | 0.615 | SMAD3 | 14.219 | FKBP1B | 5.414 | CD9 | 3.895 |
| ANK2 | 2.921 | SMAD4 | 14.593 | VSTM4 | 16.825 | FRS2 | 3.996 |
| TPM1 | 58.012 | SMAD5 | 10.406 | EGR1 | 9.129 | YPEL5 | 26.900 |
| TNNI1 | 12.452 | PRDX2 | 357.515 | SIX1 | 72.489 | BTRC | 12.350 |
| TNNT2 | 60.450 | TACC2 | 91.255 | GJA1 | 54.345 | LIPA | 40.381 |
| TCF21 | 0.000 | ADA | 54.406 | CAV2 | 17.855 | EDNRA | 22.555 |
| MYOD1 | 85.531 | JUNB | 19.555 | FGF13 | 60.769 | WNT5A | 29.052 |
| HDAC9 | 0.561 | COL8A2 | 0.258 | TEK | 3.703 | MAP2K1 | 538.223 |
